# Columbia Basin Pygmy Rabbit Recovery Planning through Structured Decision Making

**DOI:** 10.64898/2026.04.11.716011

**Authors:** Kelly Mistry, Sarah J. Converse

## Abstract

The endangered Columbia Basin pygmy rabbit (CBPR) faces multiple threats, particularly increasing risk of larger and more intense wildfires due to climate change, emerging disease, and sagebrush habitat loss due to agriculture and development. Through the use of conservation breeding, the CBPR wild population grew from a low of 16 individuals captured in 2001 to over 100 individuals in two subpopulations in 2024. However, these subpopulations are geographically proximate, with potential risk that both subpopulations could be affected by a single wildfire or disease event. Additionally, a succession of setbacks in the breeding program has prompted a natural re-evaluation point for the CBPR conservation program.

We undertook a structured decision-making (SDM) process with participants from both Washington Department of Fish and Wildlife (WDFW) and US Fish and Wildlife Service (USFWS) to develop a strategy that is sustainable and implementable for guiding management in the coming decades across the range of the CBPR, taking into account changing conditions and updated information. A population model that incorporated both demographic and high impact event uncertainties was developed to test how alternative strategies – defined by conservation breeding program, vaccination, and translocation components – affect CBPR population growth and cost objectives.

Based on analysis of the model results, we identified the following actions that appear to have the greatest potential to allow WDFW and USFWS to meet their conservation objectives for CBPR:

- Continue conservation breeding program, and possibly expand to include an island subpopulation (an isolated, unfenced area that can serve as a source for rabbits while requiring fewer management inputs)
- Continue Rabbit Hemorrhagic Disease Virus (RHDV2) annual vaccinations in both breeding and wild populations
- When juveniles are available for translocation, prioritize recovery areas that are in the establishing phase

In addition, while not analyzed explicitly in the model, discussions during the SDM process led to the identification of the following actions, which the group considered to have potential to benefit the CBPR either directly or indirectly:

- Increase the amount of suitable habitat available to pygmy rabbits
- Increase protections for existing and potential recovery areas
- Design future monitoring to better estimate survival and reproduction, with an emphasis on understanding how these vital rates vary between wild and semi-captive individuals, between vaccinated and unvaccinated individuals, and as a function of habitat factors

## 1 INTRODUCTION

The Columbia Basin Pygmy Rabbit (CBPR), a genetically distinct population of pygmy rabbits (*Brachylagus idahoensis*) found in eastern Washington, has been state and federally listed as endangered since 1993 and 2001, respectively. The CBPR faces multiple threats, particularly increasing risk of larger and more intense wildfires due to climate change, emerging disease, and sagebrush habitat loss due to agriculture and development (Gallie & Hayes 2019). Pygmy rabbits are the only native rabbit species in North America that dig their own burrows and thereby serve as ecosystem engineers in the sagebrush shrub-steppe ecosystem, benefitting plant communities by contributing to soil disturbance that increases productivity (Whitford & Kay 1999), and directly affecting vegetation structure and composition through herbivory (Huntly & Reichman 1994).

By 2001, the CBPR population in the wild had dwindled to 16 individuals due primarily to habitat loss and fragmentation (USFWS 2007, WDFW 1995), prompting a decision to capture these individuals and use them to establish a conservation breeding program. This program was successful, and individuals were released back into the wild beginning in 2007 (USFWS 2007). In 2011, the conservation breeding program transitioned from fully-captive breeding at Washington State University, the Oregon Zoo and Northwest Trek Wildlife Park to a semi-captive breeding program using permanent and semi-permanent enclosures in the CBPR home territory of the Columbia River Basin (Gallie & Hayes 2019).

The conservation breeding program has produced many individuals that have been subsequently released into wild recovery area sites, including 2,246 individuals between 2011 and 2019 (Gallie & Hayes 2019). While there have also been setbacks, two independent wild subpopulations have been established, which are currently self-sustaining as of 2024. However, these subpopulations are geographically proximate, with potential risk that both subpopulations could be affected by a single wildfire or disease event. Therefore, establishing new wild subpopulations that are more spatially distributed is a high priority for CBPR managers.

In recent years, the conservation breeding program has experienced significant challenges, resulting in relatively few juveniles produced in 2023 and 2024, and an increasing average age of breeders. A 2021 wildfire destroyed a semi-captive enclosure and its inhabitants, and in 2023, an existing semi-captive enclosure failed to produce any new breeders. This succession of setbacks in the conservation breeding program, coupled with the urgent need to establish new wild subpopulations to decrease the risk of extinction from a single catastrophic event, has resulted in a re-evaluation point for CBPR conservation.

In order to evaluate available options and make a decision about how the program should move forward over the next few decades, the management partners elected to participate in a structured decision making (SDM) process. The questions of particular interest to the partners, including Washington Department of Fish and Wildlife (WDFW) and the US Fish and Wildlife Service (USFWS), included whether to continue, reduce, or expand the current conservation breeding program, and what management actions to take to mitigate the risks associated with catastrophic events such as wildfires and emerging infectious disease.

### 1.1 Threats

#### 1.1.1 Emerging diseases

The risk of diseases that are novel to the CBPR is increasing. In the past five years, a new strain of Rabbit Hemorrhagic Disease Virus (RHDV2) has spread across much of the western United States, including parts of Oregon and Idaho in proximity to the Washington border. RHDV2 affects all species of wild and domestic rabbits, results in an extremely high risk of mortality, and is highly infectious (Bosco-Lauth et al. 2024). RHDV2-associated mortality has been recorded in pygmy rabbits, most notably in Nevada (Crowell et al. 2023). An effective vaccine does exist, and vaccinations are currently administered in the conservation breeding program and to some extent in the wild subpopulations.

However, there is uncertainty regarding whether vaccinations have a survival cost associated with the stress of capture and vaccination. The economic cost of administering vaccines is also non-negligible.

#### 1.1.2 Wildfire

A significant threat to CBPR is the increasing number, size, and severity of wildfires in the western United States due to climate change (Prichard et al. 2021, Cunningham 2024, Iglesias et al. 2022), including in the Columbia Basin. The largest recorded fires that have occurred in the Columbia Basin have occurred in the past 5 years, including a fire that burned over 200,000 acres. The Washington Department of Natural Resources collects fire data across the state, and the data for the counties in and around the CBPR-occupied area show a steadily increasing number of fires per year since the beginning of the dataset in 2008. Pygmy rabbits are not only vulnerable to direct mortality from a fire, they are also subject to increased predation and starvation risk following a fire that consumes the sagebrush they need for shelter and food (WDFW 1995).

#### 1.1.3 Loss of habitat and habitat fragmentation

Habitat loss and fragmentation, primarily due to agriculture and development, have been a primary contributor to the decline of CBPR (USFWS 2007, WDFW 1995). Pygmy rabbits are dependent on sagebrush, both as food (it makes up 99% of their winter diet) and shelter from predators (WDFW 1995). Lack of connectivity between suitable habitat patches will continue to be a challenge for recovery efforts, as lack of connectivity contributes to the demographic and genetic isolation of each wild subpopulation. Translocations of individuals between wild subpopulations may be required for the foreseeable future to maintain genetic diversity and subpopulation viability.

#### 1.1.4 Low population size

Low population size is a threat to CBPR, because a smaller population has a higher risk of extinction from natural stochasticity in vital rates as well as a loss of fitness due to lack of genetic diversity (Lande 1988). Pygmy rabbits seem to have high oscillations in demographic rates, with periodic population booms and busts (e.g. Price et al. 2010), and therefore may be particularly vulnerable to this threat (Hung et al. 2014).

In addition to the risk of individual fitness decline from loss of genetic diversity, another source of concern is the loss of Columbia Basin (CB) ancestry, which is, partially, the basis for CBPR protection as a distinct population. Management efforts in the past two decades have centered around maintaining CB genes. Between 2016 and 2019, individuals in the CBPR population were estimated to have between 9 and 54% CB ancestry (Gallie and Hayes 2019).

### 1.2 Challenges to conservation planning

The primary challenges to conservation planning raised during the course of the SDM process included the risk of extinction associated with highly uncertain, potentially highly impactful events (such as wildfire and RHDV2) and uncertainty about vital rates such as survival and reproduction, which makes predicting population responses to management actions particularly difficult. In order to address these challenges, a population model was developed that incorporates existing knowledge about demography and allows for high-impact events.

## 2 METHODS

### 2.1 Structured decision-making process

To initiate a SDM process for planning CBPR conservation, partners established a working group consisting of 10 people, 6 representing WDFW and 4 representing USFWS. The working group met for a two-day workshop in August 2024 to draft the decision problem and fundamental objectives, and to generate initial ideas for alternative conservation strategies. Through partial-and full-group follow-up meetings between September 2024 and May 2025, alternative strategies were further developed and refined, and partner input was solicited on the population and cost models (described below) that were built to evaluate the alternative strategies in terms of conservation objectives and a cost objective. A complete summary of the qualitative products of the SDM process, including a statement of the decision problem, objectives, and alternative strategies, can be found in Appendix A, with summaries below.

Several meetings were conducted between January and June 2025 to review preliminary results and revise the alternative strategies, population model, and presentation of results.

#### 2.1.1 Objectives

A variety of management objectives were identified by the working group (see Appendix A). Those that were directly evaluated in the analysis presented here include the following conservation objectives: the number of occupied wild subpopulations of the CBPR, extinction risk of the CBPR, and median total wild abundance in the CBPR. Because these three conservation objectives were highly correlated, we relied on the mean number of wild subpopulations in most comparisons of strategies.

Furthermore, we evaluated costs of management.

#### 2.1.2 Alternative strategies

The final set of alternative strategies were defined by five strategy components in three categories: two components related to the conservation breeding program, a component related to translocations, and two components related to vaccinations. Alternative strategies were constructed from all relevant combinations of options under each of these strategy components (56 total). An additional “no intervention” strategy was also evaluated; this strategy excluded all management (no breeding programs, vaccinations, or translocations), to serve as a comparison. The strategy components (and options under each) included:

- Conservation breeding program

○ Conservation breeding program component: what type of conservation breeding program will be supported. Options are semi-captive, island, both, or neither.
○ Frequency of juvenile retention component: how often are juveniles retained in the conservation breeding programs to maintain breeders in the program. Options are every year or every third year.
- Translocation

○ Wild subpopulation prioritization component: when not enough juveniles are available for translocation into all possible wild subpopulations, which subpopulations are prioritized. Options are establishing subpopulations or in-crisis subpopulations.
- Vaccination

○ Conservation breeding program vaccination component: are ongoing RHDV2 vaccinations conducted in any conservation breeding programs. Options are yes or no.
○ Wild subpopulation vaccination component: are ongoing RHDV2 vaccinations conducted in the wild subpopulations. Options are yes or no.

### 2.2 Population model

To simulate CBPR population dynamics and evaluate management strategies, we constructed a female-only, age-structured, post-breeding census population model. The modeled population consisted of juveniles (newborns) and adults (1+ years). Individuals were located in either wild subpopulations (up to six total, corresponding to current or proposed future subpopulations) or in a conservation breeding program (either semi-captive or island). Individuals were further categorized by their origin (either wild or conservation breeding). Animals could only move amongst wild subpopulations or conservation breeding programs via translocation. Individuals in the conservation breeding program were categorized as belonging to either the existing semi-captive breeding program (which may be separated into multiple sites but is assumed to be panmictic) or an island breeding program. The island breeding program is envisioned as a conservation breeding program established in a highly isolated habitat fragment with natural barriers that would restrict movement of individuals in or out.

Population sizes for the currently occupied wild subpopulations (Sagebrush Flats and Beezley Hill) were initialized based on the estimated subpopulation size in 2024 (∼68 individuals in each, per WDFW scat surveys in winter 2024), assuming a sex ratio of 1:1. We used an initial age distribution determined by the stable age distribution calculated from each simulation’s survival parameters and reproduction parameters drawn from a Poisson distribution with a mean describing the mean number of kits per female (7.27; calculated from data in DeMay et al. 2016). The initial population in the semi-captive breeding program was based on the known number of females (5) in the enclosures as of December 2024, classified as adults in year 1 of the model, which starts in March 2025.

We incorporated parametric uncertainty and stochasticity throughout the population model, to account for lack of information and for the high variability in pygmy rabbit population dynamics. To incorporate parametric uncertainty about RHDV2 outbreaks, 100 values were drawn from parameter distributions, constructed via expert elicitation, for the following parameters: the annual risk of an RHDV2 outbreak occurring in the pygmy rabbit population, the probability of early detection given that an outbreak occurs, and the annual mortality from vaccine-related capture myopathy. To incorporate uncertainty about demographic parameters, 100 values for survival coefficients from a mark-recapture model for survival (described below), as well as the standard deviation of a random effect of day-of-year in the survival model, were drawn from posterior distributions. We therefore had 100 values for each parameter in the model. Each of these sets of values were used for 100 simulations, resulting in 10,000 simulations under each strategy. In each simulation, the population was simulated forward for 20 years.

Annual stochasticity was included in each of the 20 years for several processes (annual survival, reproduction, sex of offspring, start years for establishing new recovery areas, RHDV2 outbreak occurrence and timing, and wildfire occurrence and size) as described below.

To evaluate performance of strategies, we calculated mean values across all 10,000 simulations for a given strategy, with means calculated for the number of extant wild subpopulations, whether the population overall was extant at that time interval (i.e., a probability), and the median total wild abundance. To quantify parametric uncertainty, we calculated mean values for these three metrics across the 100 simulations for each of the 100 parametric sets and calculated the upper and lower 95% quantiles of these mean values across the parametric sets. All model outputs were calculated at years 5, 10, 15, and 20 in each simulation to compare strategy performance through time.

To evaluate the impact of wildfire and RHDV2 on outcomes, the model was run for all strategies under four variations: with both wildfires and RHDV2, with only wildfires (i.e., probability of RHDV2 outbreak set to 0), with only RHDV2 (i.e., probability of wildfire set to 0), and with neither RHDV2 nor wildfires.

#### 2.2.1 Annual survival

Monitoring data collected since 2016 in the conservation breeding program and in the wild subpopulations were used to conduct a mark-recapture analysis to estimate survival, with survival modeled as a function of age (juvenile or adult) and location type (conservation breeding program or wild subpopulation). Due to challenges with estimating survival (i.e., juvenile survival estimates appeared to be unrealistically high, see Appendix C for details), we did not use these estimates in the population model. Thus, only the estimates of adult survival from this analysis were used in the population model.

Juvenile survival was set to 75% of adult survival in all cases. No empirical information is available to directly compare juvenile and adult survival, and the value of 75% was chosen based only on basic considerations of life history theory, and with the expectation that the strategy outcomes were unlikely to be directly affected by this parameter choice. Further research on survival to better elucidate relative survival of adults and juveniles, or to better estimate juvenile survival on its own, would be a helpful addition to a future version of this model to either update the value for relative survival or provide a direct estimate of juvenile survival.

Survival of adults in each location type *s* (conservation breeding program or wild subpopulation) in each year *y* of each simulation *j* (𝜙_age=2,*s,y,i*_) was calculated using the simulation-specific coefficient values (intercept, age, and location-type coefficients: 𝛼*_int_*[𝑗], 𝛼_age=2_[𝑗], and 𝛼_s_[𝑗] respectively) drawn from parametric distributions. In addition, in the mark-recapture analysis we modeled survival on a daily scale with a random effect of day-of-year. Thus we sampled from the parametric distribution for the standard deviation of the random effect (𝜎[𝑗]). Each year, we then randomly sampled *i* = 365 deviates, as:

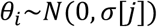

and then took the product of 365 daily survival estimates to calculate annual survival, as:

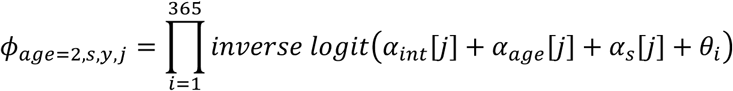

For the strategies that included wild vaccination, an additional source of annual mortality was subtracted from adult and juvenile wild survival to account for the possible survival cost of administering vaccinations. There is virtually zero mortality from complications related to the vaccine itself (K. Haman, Washington Department of Fish and Wildlife, personal comm.), but for individuals in the wild, there may be a small amount of mortality associated with capture myopathy from the additional captures required to administer vaccinations. The probability distribution for this added mortality parameter was developed through expert elicitation (see Appendix B).

#### 2.2.2 Annual reproduction

DeMay et al (2016) recorded the number of juveniles produced by each female in the conservation breeding program in each year from 2012 to 2014, ranging from 0 to 33 juveniles per female per year (Figure 1). For each adult female in each year, a value is drawn from the specific set of values recorded by DeMay et al. (2016) representing the number of juveniles produced by that female. The sum of these draws across females is the total number of juveniles, and a binomial draw from this number, with probability 0.5, provides the number of juvenile females.

**Figure 1.**
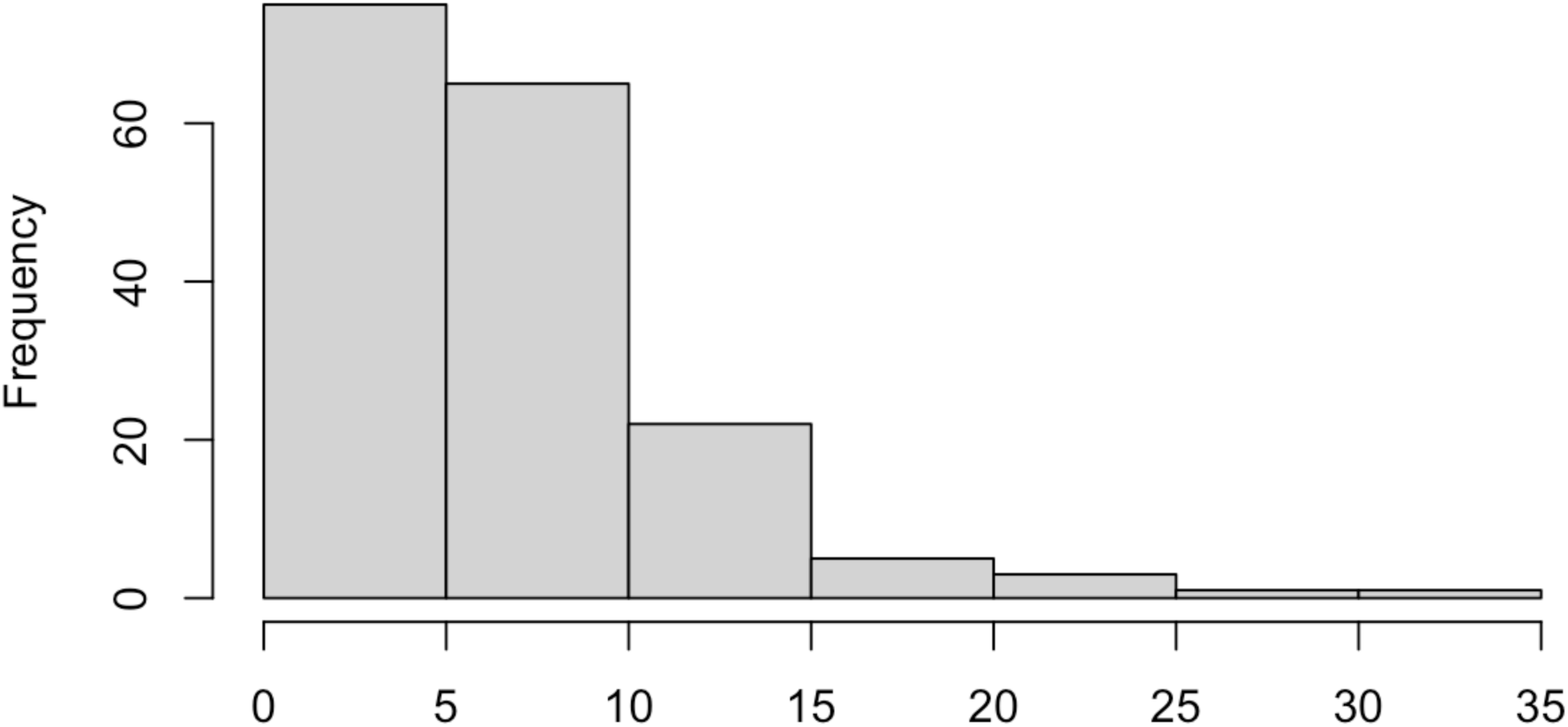
The number of kits per female per year recorded by DeMay et al. 2016. Data were collected between 2012 and 2014 in the semi-captive breeding program in Washington.

#### 2.2.3 Density dependence

To prevent the number of rabbits in wild subpopulations or conservation breeding programs from growing to improbable values, an upper limit for abundance (𝑁*_max_*) was determined. For wild subpopulations and the island breeding program, this upper limit was calculated based on the area of the site in hectares (𝐴) and a maximum per-hectare density. The maximum per-hectare density was calculated using a non-linear relationship between abundance and burrow density estimated by Price and Rachlow (2011; 𝑎 = 0.2508 and 𝑏 = 0.1564) and the maximum burrow density from an unpublished dataset provided by J Rachlow (University of Idaho, unpublished; 𝑥_3_ = 2.61), collected from two wild populations in Idaho over a 12-year period:

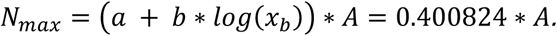

For the semi-captive breeding program, the upper limit for abundance was set to 4000 individuals (distributed across up to four enclosures), based loosely on the highest annual yield in the past (over 800 juveniles in 2014) and assuming that approximately 40% of a population is juveniles immediately after the breeding season (i.e., 2000 individuals) and then doubling that number to be conservative. When a wild site or the island site exceeded its abundance limit, abundance was reduced to the upper limit value, thereby capping the population.

Density dependence also has a role in determining which wild subpopulations are eligible to receive translocated juveniles in any given year. If the wild subpopulation reached over 10% of its maximum abundance in any year, then it was considered ineligible to receive translocated juveniles. This rule was based on current management practice, where translocations occur in relatively small release sites (each is 1 acre, fenced) established for 6 to 8 months inside each recovery area to facilitate soft releases. If the area inside and immediately around these release sites becomes too densely occupied, then releasing more individuals in the same area may be detrimental to their survival.

#### 2.2.4 Wild recovery area sites

The model included two occupied recovery areas (RA; Sagebrush Flats and Beezley Hills) and four proposed recovery areas (Figure 2). The proposed RA sites were selected through a comprehensive review process conducted by WDFW and USFWS, which identified sites most appropriate to prioritize for translocations, based on habitat suitability and the relative ease with which conservation benefit agreements (CBAs) could be developed with landowners (Gallie et al. 2024).

**Figure 2.**
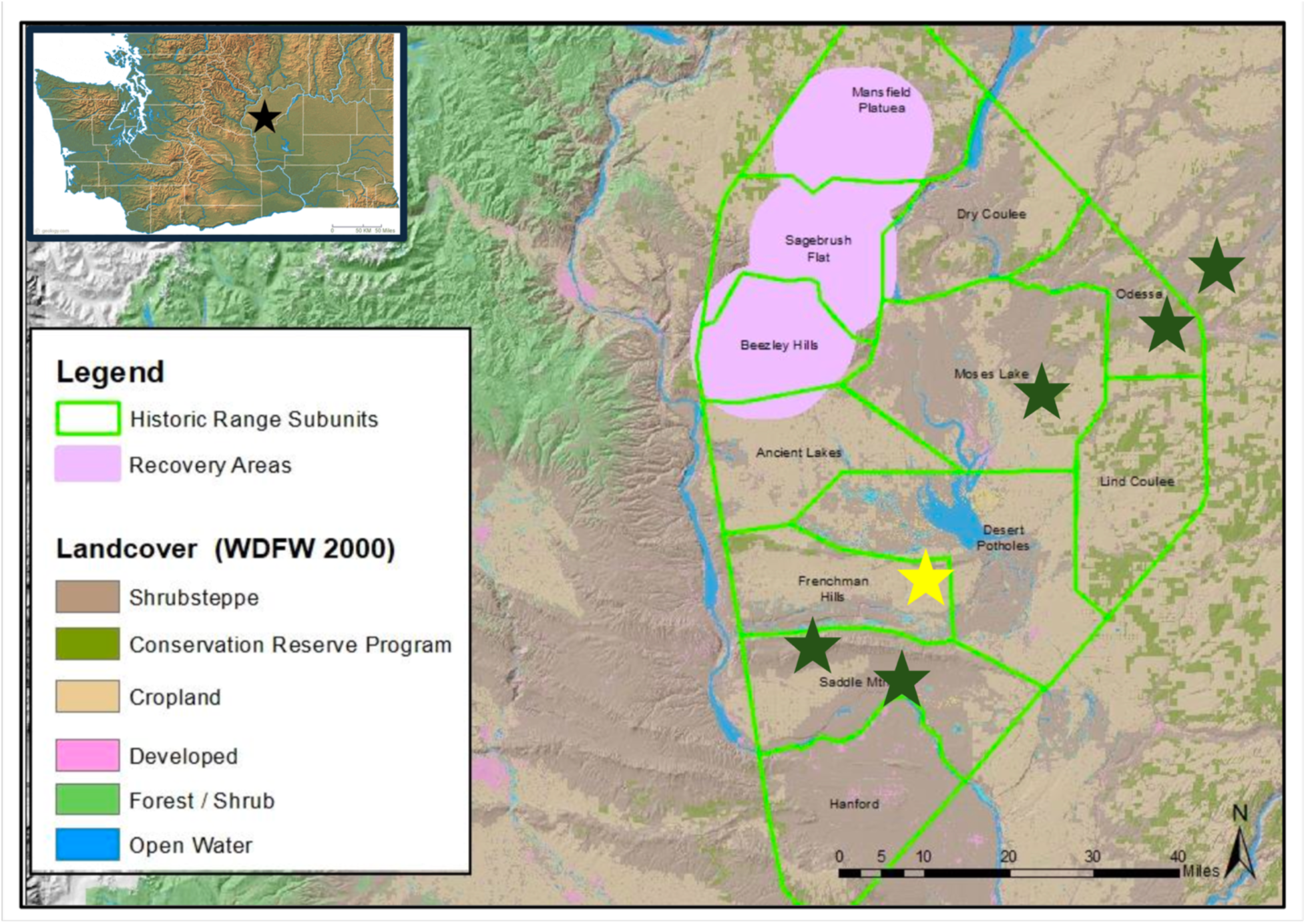
Map of the existing and proposed wild recovery areas. The existing recovery areas, Sagebrush Flats and Beezley Hills, are shown in pink (Mansfield Plateau was a recovery area but was burned in 2019 and is no longer viable habitat for pygmy rabbits) within the green outer line showing the borders of the area defining the historical range of the CBPR distinct population segment (DPS), as defined by USFWS in the endangered species listing. The green stars mark the proposed recovery areas, and the yellow star delineates a suitable location for a natural “island” breeding program in Frenchman Hills. This figure was originally created in Gallie et al. 2024.

There is uncertainty regarding how long it will take to secure CBAs and inter-agency agreements (for government-owned sites) to prepare the proposed RAs (Blackrock Coulee, Lakeview, Saddle Mountain, and a private ranch referred to here as Ranch) to receive pygmy rabbits, as each site differs in the number and types (e.g., government, private) of landowners. Timeline ranges were developed for each proposed RA based on expert judgment from state and federal managers who have acquired CBAs for pygmy rabbits in the past, specifying the minimum and maximum number of years before the site could be made available for translocations. In addition, there is uncertainty regarding how much effort can be applied to developing these sites, i.e., how many sites can be in development at one time, based on agency workloads and competing priorities. Both of these sources of uncertainty were used to develop a start year for each site in each simulation; this start year was the first year that the site was eligible to receive translocated individuals. In each simulation, the number of sites that could be in development in each year (site rate) was determined by drawing a value of 1, 2, or 3 (with equal probability). Then, each site’s development timeline was determined by drawing a value between the site’s minimum and maximum timeline range. The site development order was random, with the caveat that two sites under the jurisdiction of the same WDFW private lands biologist could not both be in development at the same time (only relevant if site rate was 2 or 3 per year). The start year of each site for each simulation was then determined based on the order of the sites, the site rate, and each site’s timeline. For example, if the site order was Blackrock Coulee, Lakeview, and Saddle Mountain, the site rate was 2, and the site timelines were 2, 5, 3, and 2 respectively, then the start years for the first set of sites would be year 2 for Blackrock Coulee, year 5 for Ranch. The second set of sites would have start years of year 5 for Lakeview and year 7 for Saddle Mountain, as each of these sites would enter the development phase once one of the first sites was developed.

In the model, the subpopulation of each RA site was evaluated each year to determine what, if any, translocation actions could occur in that year. Sites were defined as “establishing” until they had been occupied continuously for five years after becoming available for translocations, based on the simulated start year for the site. Because abundance can vary quite significantly from year to year due to interannual variation in survival, continuous occupation for at least 5 years regardless of abundance is sufficient, in the context of the model, for considering a site to be established. The existing sites, Sagebrush Flats and Beezley Hills, were considered established in year 1. If the population was extirpated from an established site, then it was re-classified as establishing in the next year. The exception to this was if the site’s subpopulation died out due to wildfire, in which case the site was no longer available for translocations for the rest of the management time horizon (20 years), as sagebrush habitat takes decades to recover from a wildfire. Established sites were further evaluated for whether they were declining (negative growth between year *t*-1 and year *t*) or in crisis (fewer than five females left in the subpopulation in year *t*). Sites that were categorized as establishing or in-crisis were prioritized as recipients for translocation of available juveniles, according to specific decision rules described below.

#### 2.2.5 Conservation breeding programs

There are two strategy components relevant to the conservation breeding programs: the type of conservation breeding program(s) and the juvenile retention schedule in the conservation breeding program. The semi-captive program using mobile enclosures is currently in existence, while the island program has not been attempted previously, but is here conceived as using a proposed recovery area site (Frenchman Hills) as an isolated “island.” This site was identified for the island program because it is in a valley bordered by either water or cliffs on all sides, making both emigration and immigration very difficult. The semi-captive breeding program has been operating in its current form since 2017 and consists of up to four mobile enclosures that are moved to new sites every three to five years. Historically, two to three enclosures have typically been in operation at any given time. In the model, all enclosures were treated as one population, and it was assumed that two enclosures would be in operation while the program was operating.

There is some uncertainty about the logistical difficulty of establishing an island breeding program at the Frenchman Hills site. In the model, for any strategies with an island breeding program, this uncertainty was represented by a range of possible start years (1 to 3), from which we sampled for each simulation. For strategies with only an island breeding program, the existing semi-captive breeding program continued to operate until the island breeding program was ready to receive pygmy rabbits. At that point, all individuals were translocated from the semi-captive breeding program to the island breeding program. For strategies with both programs, the semi-captive breeding program continued to operate alongside the island breeding program in the model. For strategies with neither breeding program, existing individuals in the semi-captive breeding program were translocated into the Sagebrush Flats RA in year 1.

Similar to the wild subpopulations, any conservation breeding programs were evaluated in the model in each year to determine what translocations, if any, could occur in that year. An island breeding program was classified as establishing for the first three years after the island site became available, after which it was classified as established as long as it was occupied. During the establishment phase, up to 15 juveniles produced in the island breeding program were retained to allow for growth, but if more than 15 juveniles were produced, then the remainder were available for translocation. The semi-captive breeding program was considered established in year one and all juveniles produced in the program were available for translocation unless juveniles were to maintain the program.

In each year that juveniles were retained to maintain a conservation breeding program, between three and five juveniles were retained and any remainder were available for translocation. If the strategy option for juvenile retention was every three years, then juveniles were retained on this schedule beginning in year 3 for the semi-captive program, and in the third year after the start year for the island program. Either conservation breeding program was eligible for “rescue,” i.e., was an eligible recipient of translocated individuals, if it had two or fewer females and was classified as established.

There were several conditions that resulted in the deliberate closure of the semi-captive breeding program, in which case all individuals in it were translocated to a wild subpopulation. The semi-captive breeding program could close if wild subpopulations were established in at least four different fire zones (fire zones are described in ***2.2.8 Wildfire risk***), or if an island breeding program was established and there were wild subpopulations in at least 3 different fire zones. It was assumed that the island breeding program would not deliberately be closed under any condition if it existed.

#### 2.2.6 Translocations

There is one strategy component related to translocations; which types of wild sites are prioritized to receive translocations: establishing or in-crisis sites. Establishing was defined as above (sites that have not been occupied continuously from year 𝑡 − 5 to year 𝑡) and in-crisis was defined as a site with a subpopulation with fewer than five females in year *t*.

Retention of juveniles in a conservation breeding program, as described above, was prioritized before the program was sourced for translocations. After that priority was met, the possible types of translocations that could occur (in order of priority if more than one translocation was possible in year *t*) were: establishing the island breeding program, translocations to the semi-captive or island breeding program if the program was a candidate for rescue, periodic translocations to the semi-captive or island breeding program to bolster genetics, annual translocations of available juveniles from the semi-captive or island breeding programs to wild subpopulations (with the site priority set according to the strategy), and translocations between wild subpopulations if the breeding program(s) did not exist or were not producing enough juveniles for all of a given strategy’s priority wild subpopulations (with the site priority set according to the strategy). The number of individuals translocated under any of these types was based on the number of available juveniles in each year and the type of translocation. Rules guiding each of these translocation types are described below.

The establishment period for an island breeding program included the first three years after the island program’s start year, and the number of juveniles translocated into it in each model year was randomly selected from between 12 and 18 juveniles. The range of values for number of juveniles to translocate in per year was based on expert advice about the minimum number of individuals that had been used in the past to successfully start the breeding program (18 individuals in 2001; USFWS 2007). The semi-captive breeding program was the preferred source, if it existed and had available juveniles. If it did not have enough available juveniles, then available wild juveniles could be translocated in as well.

A conservation breeding program was eligible for rescue if there were 2 or fewer individuals left in the population. Sources for rescue in order of preference were the other conservation breeding program if it existed, then wild sources. The number of juveniles for this type of translocation was randomly drawn between 5 and 8.

Periodic translocations into existing breeding programs for the purposes of maintaining genetic diversity first drew from the other breeding program’s available juveniles, if another program existed. Otherwise, wild subpopulations were considered. The number of juveniles for this type of translocation was randomly drawn between 3 and 5. Genetic translocations occurred as often as every 3 years into each breeding program if juveniles were available; for the semi-captive breeding program they started in year three, and for the island breeding program they started three years after the start year in a given simulation.

Translocations into wild subpopulations represent a strategy component. Once any uses for juveniles described above were met, remaining juveniles from the conservation breeding program were translocated into priority wild subpopulations in the model. The priority subpopulations were either establishing sites or in-crisis sites. If there were sites that fit the priority criteria, then available juveniles were equally split between these sites, with a minimum of 3 juveniles translocated in each group. If there were no priority sites, then juveniles were evenly split between wild sites that were eligible to receive juveniles. The only condition that made a wild site ineligible to receive juveniles was if it had reached over 10% of its maximum abundance (see ***2.2.3 Density dependence***).

In the absence of a breeding program or if there were not enough available juveniles from the breeding program to accommodate all priority wild subpopulations, then wild subpopulations could be used for wild-to-wild translocations. For wild-to-wild translocations, a value was randomly drawn between 5 and 8 for each priority recipient site. Wild sites were eligible to be sources if the site had had positive growth for at least the past 2 years and it had more than 40 females. A maximum number of juveniles that could be sourced from a wild subpopulation was set at 10% of the subpopulation abundance.

#### 2.2.7 RHDV2 and vaccinations

Rabbit hemorrhagic disease virus type 2 (RHDV2) is a highly infectious virus posing a high mortality risk to pygmy rabbits. This virus was first detected in Europe in 2010 and first appeared in the United States in 2018 (Asin et al. 2021). There have been no major outbreaks in Washington state as of June 2025. It is challenging to estimate the annual probability of an outbreak occurring or the probability of advanced detection if an outbreak were to occur. An effective vaccine that greatly increases survival after exposure exists and currently is administered to all individuals translocated out of the conservation breeding program annually, as well as the majority of individuals in the conservation breeding program during routine captures. Periodic vaccinations occur in the wild subpopulations as well, administered to an estimated 30 to 50% of each subpopulation. The vaccine requires two doses, an initial shot followed three weeks later by a booster, and full immunity occurs two weeks after the booster dose is administered (five weeks after the initial shot).

Two strategy components exist regarding vaccinations for RHDV2, including whether there is an ongoing (annual) vaccination program for individuals in the conservation breeding program, and whether there is an ongoing vaccination program for individuals in the wild subpopulations. Regardless, emergency vaccinations would occur for both groups if an imminent outbreak was detected, as described below.

Three parameters relevant to RHDV2 were elicited from experts (see methods, Appendix B).

Expert elicitation was conducted to estimate the annual probability of an RHDV2 outbreak occurring in the CBPR; the probability that, if an outbreak occurs, it will be detected soon enough to allow partners to mount an emergency vaccination effort (detection within a 100 mile radius of CBPR would trigger an emergency vaccination effort); and the added annual mortality from capture myopathy during vaccinations in wild subpopulations.

If annual vaccination in the conservation breeding program(s) was part of a strategy, then we assumed that all individuals were vaccinated annually. If annual vaccination in the wild population was part of the strategy, then a binomial draw with index equal to the number of rabbits in each wild subpopulation determined how many individuals were vaccinated in each year, with probability = 0.3 (∼30% of each subpopulation). This probability was a conservative estimate of how much of the total population was likely to be vaccinated using current practices.

The vaccinated portion of each wild subpopulation was subject to an adjusted annual survival; we drew a value for annual mortality from capture myopathy from the parameter distribution for each simulation and subtracted it from each year’s annual survival. This adjusted annual survival was used for the portion of each wild subpopulation that was vaccinated in each year.

In each year, a Bernoulli draw with the simulation-specific annual outbreak probability determined whether an RHDV2 outbreak occurred in that year (1 = outbreak, 0 = no outbreak). If an outbreak occurred, another Bernoulli draw with the simulation-specific probability of advanced detection of the outbreak determined whether an emergency vaccination process was mounted (1 = detected, 0 = not detected). Both of these probability distributions were constructed using an expert elicitation (see Appendix B for elicitation description).

If an outbreak occurred, then the entire population (both wild subpopulations and the conservation breeding programs) was subject to mortality, with the risk of mortality based on vaccination status. Bosco-Lauth et al. (2024) found that the survival probability for unvaccinated individuals exposed to the disease was 0.05, while all individuals in their study (55 New Zealand white rabbits) vaccinated with two doses of the vaccine survived. We recognize that for vaccinating individuals in wild subpopulations, some individuals will not be captured and will receive no vaccine doses, and some will only be captured once and will receive one dose. These individuals will not receive the full protection from the vaccine, although there is anecdotal evidence that receiving a single dose does provide some protection. Based on the judgment of the WDFW wildlife veterinarian (K Haman, Washington Department of Fish and Wildlife, personal comm.), survival for vaccinated individuals during an outbreak was set at 80% in the model. Additional mortality was additive to baseline annual mortality for unvaccinated rabbits and vaccinated rabbits in the conservation breeding program and to the adjusted annual mortality for vaccinated wild rabbits.

If an outbreak was detected before it occurred, emergency vaccinations occurred, with stochasticity incorporated to represent variability in the amount of advance notice. First, we calculated the number of days of the emergency vaccination effort based on the time between detection and the outbreak reaching the CBPR. The animals vaccinated in that period were assumed to be fully and effectively vaccinated, implying a minimum amount of time between an outbreak detection and the virus reaching the pygmy rabbit population of 5 weeks. Based on anecdotal evidence (K. Haman, Washington Department of Fish and Wildlife, personal comm.), 3 months was used as the maximum amount of advance notice, based on the speed at which outbreaks in nearby states have taken to travel approximately 100 miles. The number of vaccination days was the amount of time between this minimum and maximum, randomly sampled from 2 to 49 days.

For each day available for vaccination, a Poisson draw was used to select the number of individuals vaccinated on that day, with an expectation of 5 individuals per day. The sum of these daily vaccinations was the maximum number of individuals that could be vaccinated in the emergency effort. These vaccinated individuals were then attributed to the existing wild subpopulations in random order, with a limit of a maximum of 60% of any given wild subpopulation vaccinated. The newly vaccinated individuals in each wild subpopulation were randomly selected, in terms of age and previous vaccination status.

#### 2.2.8 Wildfire risk

We modeled wildfire impacts on both wild subpopulations and conservation breeding programs within the CBPR recovery area. Substantial uncertainty and stochasticity are challenges for modeling wildfire impacts.

We began by estimating the number of wildfire ignitions occurring annually inside the historical range of CBPR (see Figure 2). We used annual fire data collected by the Washington Department of Natural Resources from 2008 to 2024 (data subset to the 5 counties in and around the historical range; Grant, Douglas, Franklin, Benton, and Adams counties) to construct a Poisson distribution for the number of fires per year. We estimated an annual mean of 9.8 ignitions. In each year of each simulation, we used this distribution to draw the number of ignitions occurring anywhere in the historical range in that year.

Each fire was then assigned an ignition location by randomly selecting the index of a cell from a 5-mile by 5-mile grid placed over the historical range and the immediate surrounding area (Figure 3). The current and proposed wild subpopulation sites and the island breeding program site were assigned grid locations based on their spatial location, with each site occurring in only one grid cell. Each wildfire ignition was then randomly assigned a size class of 0, 1, 2, or 3. The probability of a fire being a member of a given size class was based on the frequency of fires in that size class in the DNR data set (see Table 1 for size class descriptions).

**Figure 3.**
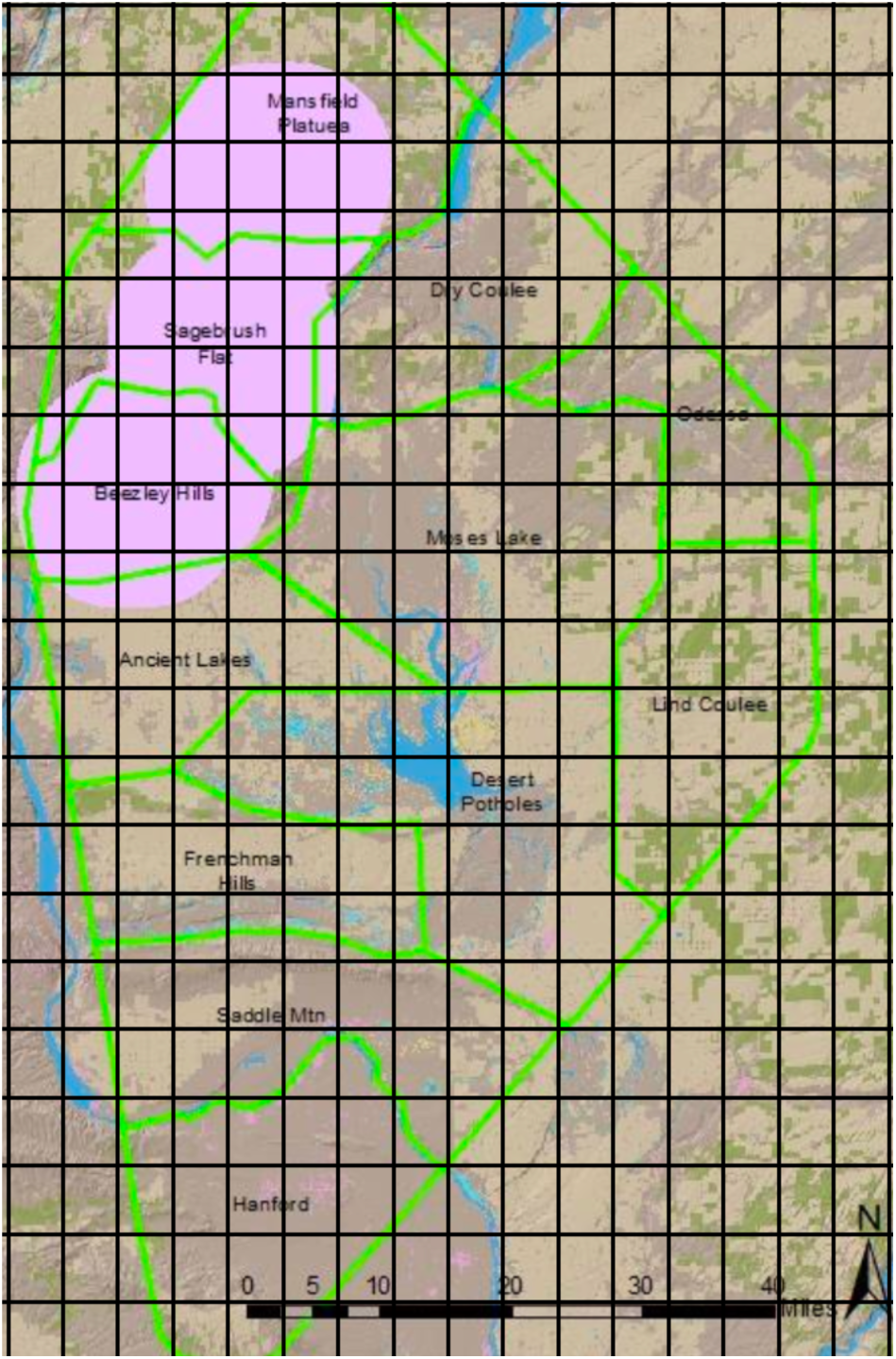
Map of the historical range of the CBPR with overlaid grid. The green outer boundary delineates the historical range, and the black lines show the 5 x 5 mile grid used to spatially distribute wildfire ignitions. The pink areas are where past and current pygmy rabbit subpopulations are located (Sagebrush Flats and Beezley Hills are the currently existing subpopulations). The proposed locations of the new wild subpopulations occur within the interior green-bordered regions labeled Odessa, Moses Lake, Frenchman Hills, and Saddle Mtn. The figure underlying the grid was created by Gallie et al. 2024.

**Table 1.** The name of each fire size class, with the size range represented by that class, the probability of assigning the size class to a fire, and the number of 5 x 5 mile grid cells that the size class will potentially affect.

| Size class | Size range (in square miles) | Probability | Grid cells affected |
| --- | --- | --- | --- |
| Zero | $\leq 0.1$ | 0.299 | 0 |
| One | 0.1 to 25 | 0.689 | Ignition cell only |
| Two | 25 to 225 | 0.006 | Ignition cell and potentially all surrounding cells (9) |
| Three | $\geq 225$ | 0.006 | Ignition cell and if it occurs inside a fire zone, then potentially all other cells in the fire zone |

The impact of wildfires on the pygmy rabbit population was determined by the size and spatial location of the fires in each year. A fire with a size class of 0 did not affect any subpopulation. A fire with a size class of 1 affected any wild subpopulation site or island breeding program site if the ignition location was in the same grid cell as the site, with 100% mortality for pygmy rabbits at that site. A fire with a size class of 2 affected a wild subpopulation or island breeding program site if the site was located either in the ignition grid cell or in any of the eight grid cells surrounding the ignition location, with 100% mortality. Fires of size class 3 affected a wild subpopulation or island breeding program site if the ignition location grid cell was in the same fire zone as the site grid cell, with 100% mortality. Fire zones were delineated areas where wildfires would be expected to spread easily, with natural barriers between zones including rivers, irrigated agriculture, and towns. These zones were drawn by WDFW biologists using a satellite map and their knowledge of the area (Figure 4) and served as a way to model spread of very large fires.

**Figure 4.**
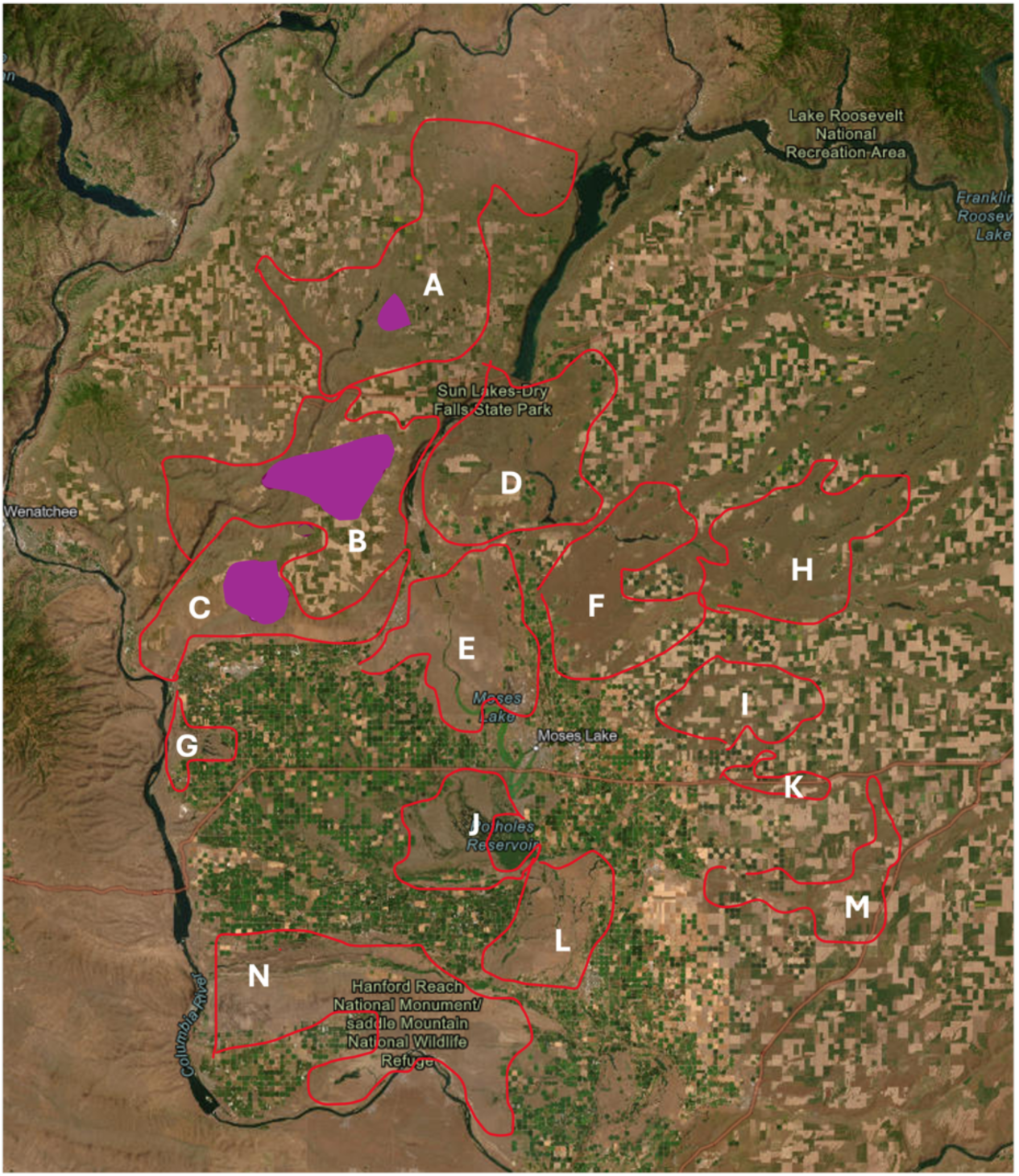
Satellite map of the historical range of CBPR and the surrounding area with overlaid fire zones. Fire zones were drawn directly on this map by WDFW biologists, delineated by red lines and labeled with white letters. The purple polygons in the graphic indicate locations where rabbits have been captured in the past in the two existing recovery areas, Sagebrush Flats (A and B) and Beezley Hills (C). Underlying map was generated by WA Dept of Fish & Wildlife biologist.

For any wild subpopulation or island breeding site affected by a fire, the site was removed from the list of sites eligible for future translocation for the remainder of that simulation. It takes the sagebrush ecosystem around 20 years to recover sufficiently from a wildfire to support pygmy rabbits, which is a timeframe beyond the scope of this model simulation.

For the semi-captive breeding program, the model assumed that there were 2 mobile enclosures operating at any given time if the program was active, and their spatial location was not fixed. Therefore, Bernoulli draws were used to determine whether any fires affected each semi-captive enclosure in each year, with the probability calculated using the number of fires that occurred in that year divided by the total number of grid cells (336). If one enclosure was affected by a wildfire, then the semi-captive breeding program experienced 50% mortality, and if both were affected, mortality was 100%. However, the breeding program was eligible to continue to operate in the next year, if there were survivors or available juveniles from wild or island sources to reestablish it.

### 2.3 Reduced wild survival scenario

The coefficient estimates for location type from the survival analysis indicated that the mean survival for animals in the wild population was higher than in the conservation breeding program (Figure 5). This was a surprising result and may be an artifact of the dataset (see Appendix C). This result was also a likely driver for the high wild abundances that many of the simulations achieved with or without a breeding program, and the relatively low extinction risks for all strategies compared to earlier versions of the model that used static annual survival probabilities (based on literature review), where wild survival was lower than in the conservation breeding program for both ages. As a sensitivity analysis, we ran all strategies for all simulations of the model with the posterior distribution of the wild location type coefficient restricted to negative values only, thereby guaranteeing that wild individuals would have generally lower survival than breeding program individuals (Figure 6). We refer to this version as the reduced wild survival scenario.

**Figure 5.**
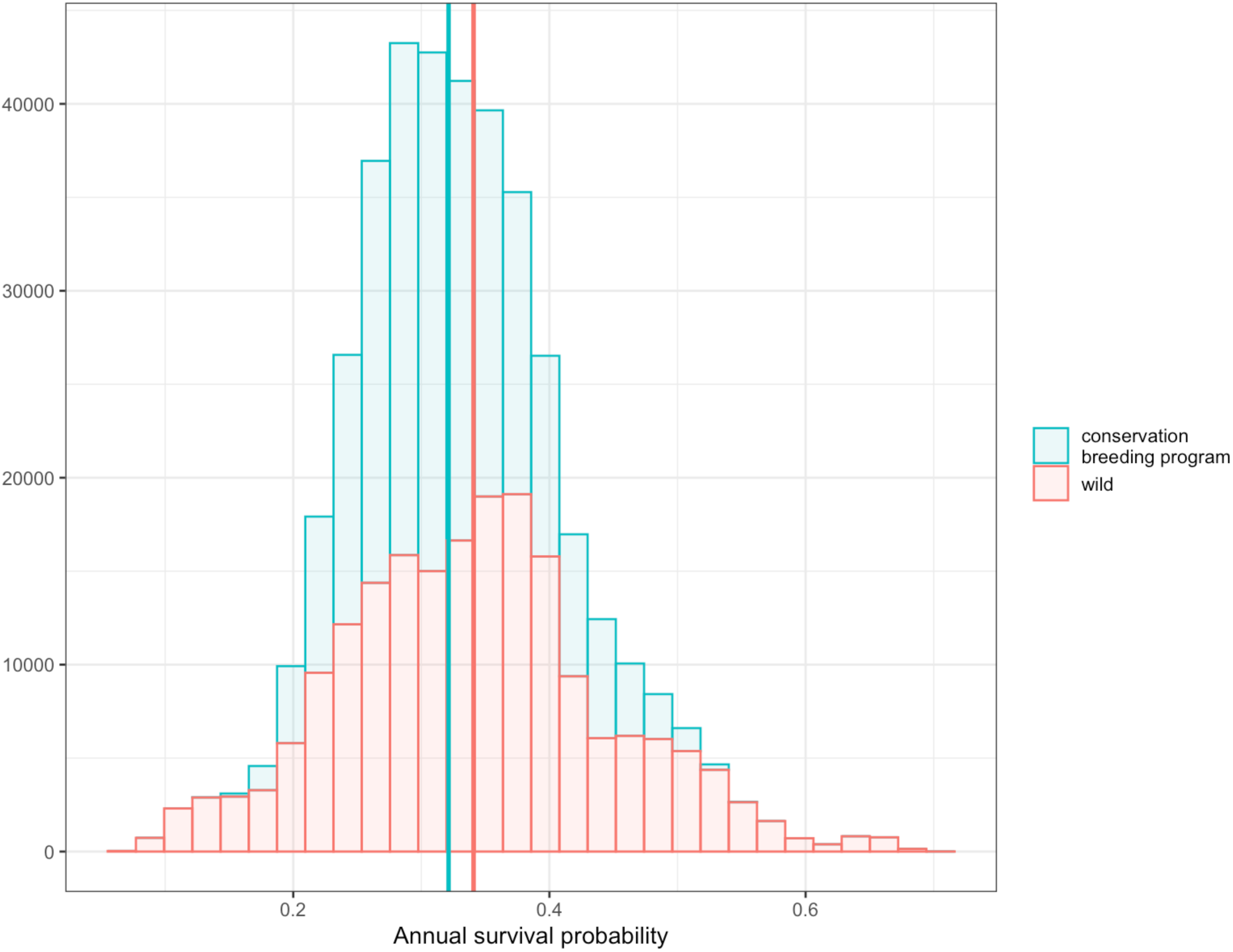
The adult survival probabilities produced by simulations in the full version of the model, with all coefficients drawn from their entire posterior distributions from the survival analysis. Wild survival is shown in red and conservation breeding program survival is shown in blue. Mean survival probability values for each location type are shown with vertical lines, red for wild (0.34) and blue for conservation breeding program (0.32).

**Figure 6.**
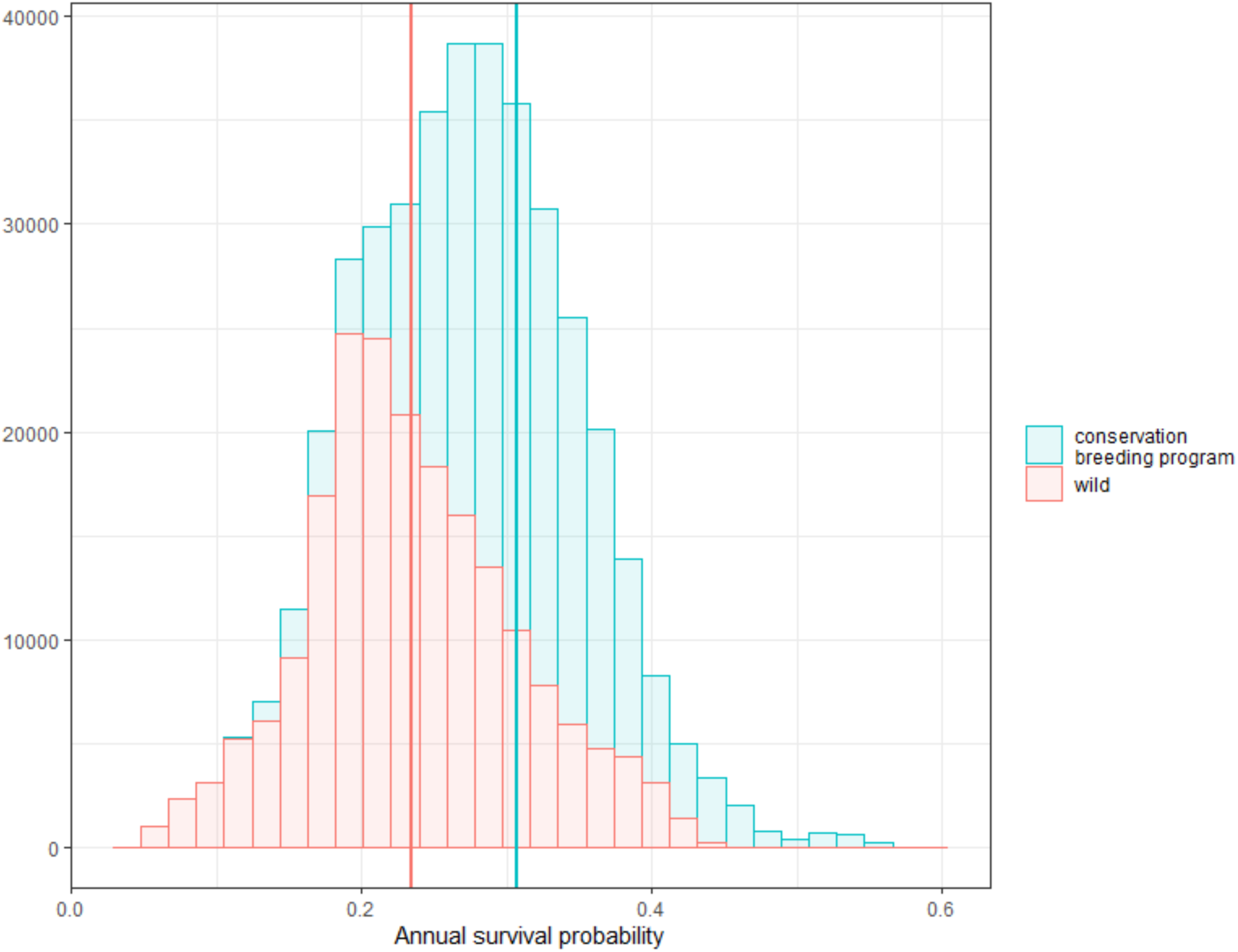
The adult survival probabilities produced by simulations in the reduced wild survival scenario. Mean survival probability values are shown with vertical lines, red for wild (0.23) and blue for conservation breeding program (0.31).

### 2.4 Cost model

We built a model to estimate costs of management, which draws on both strategy characteristics and population characteristics (specifically, the number of individuals in the population) to determine costs. All strategies were subject to a base cost, which covered up to 3 staff members, equipment maintenance, and routine replacements, based on the budget allocated by WDFW for pygmy rabbits in 2024 ($285,000). Strategies that incorporated vaccinations also incurred costs related to the vaccine ($20 per rabbit vaccinated per year, based on 2024 pricing). Additionally, strategies that included the semi-captive breeding program were subject to an annual probability that enclosure fences would need to be replaced, primarily due to wildfire. This probability was based on three wildfire events that required enclosure fence replacements in a 14-year period (i.e., 3/14 = 0.21). The cost of each enclosure fence replacement was drawn from a uniform range ($15,000 to $20,000). Costs were calculated for each simulation, then averaged across simulations to produce mean cost estimates for each strategy.

### 2.5 Evaluation of alternative strategies

Alternative strategy results were plotted on two dimensions: the mean number of extant wild subpopulations and the mean cost. Uncertainty was included in plots. This provides a visualization of the Pareto frontier (Converse 2020). Alternative strategies on the Pareto frontier are those for which it is not possible to gain performance on one objective without giving up performance on another objective. In other words, the Pareto frontier (also referred to as the set of Pareto-efficient strategies) is composed of the cheapest strategy for a given number of extant wild populations and the largest number of extant wild populations for a given cost. We evaluated both overall strategies and individual strategy components across all strategies in intermittent evaluation years (years 5, 10, 15, 20) in this way. Strategy components were also evaluated by calculating the difference in the mean number of wild subpopulations and mean cost for paired strategies, where all other components are identical except the component being compared. The distributions of these differences were described using the mean and the range between the minimum and maximum values.

## 3 RESULTS

### 3.1 Strategy evaluation

Overall, there were five strategies that produced the highest three values for mean number of wild subpopulations in multiple evaluation years (including year 20 for four out of the five) and for multiple model variant conditions. Of these strategies, all of them had at least one breeding program (three had a semi-captive program, one had an island program, one had both breeding programs), and all prioritized establishing wild sites over in-crisis sites when allocating individuals for translocations. There was more variety in the vaccination options of these strategies, as well as the juvenile retention schedule options for the breeding program. However, for model variant conditions that included RHDV2 (RHDV2 only or both wildfire and RHDV), vaccination was key to achieving population objectives. Of these RHDV2 variants, nine strategies produced the top three values for the mean number of wild subpopulations for multiple evaluations years. Out of those nine, 77.8% (7 strategies) vaccinated the conservation breeding program population, 66.7% (6 strategies) vaccinated the wild populations, and 55.6% (5 strategies) vaccinated both the conservation breeding program and the wild populations. Only one of these strategies had no vaccination in either type of population.

Four strategies had the highest mean number of wild subpopulations (by the end of the study as well as earlier) for multiple model variant conditions. The mean number of wild subpopulations in these four strategies ranged from 2.06 to 3.30 wild subpopulations depending on the model variant conditions (the highest values occurred with no RHDV2 outbreaks, the lowest values occurred with both RHDV2 and wildfire; see Figure 7).

**Figure 7.**
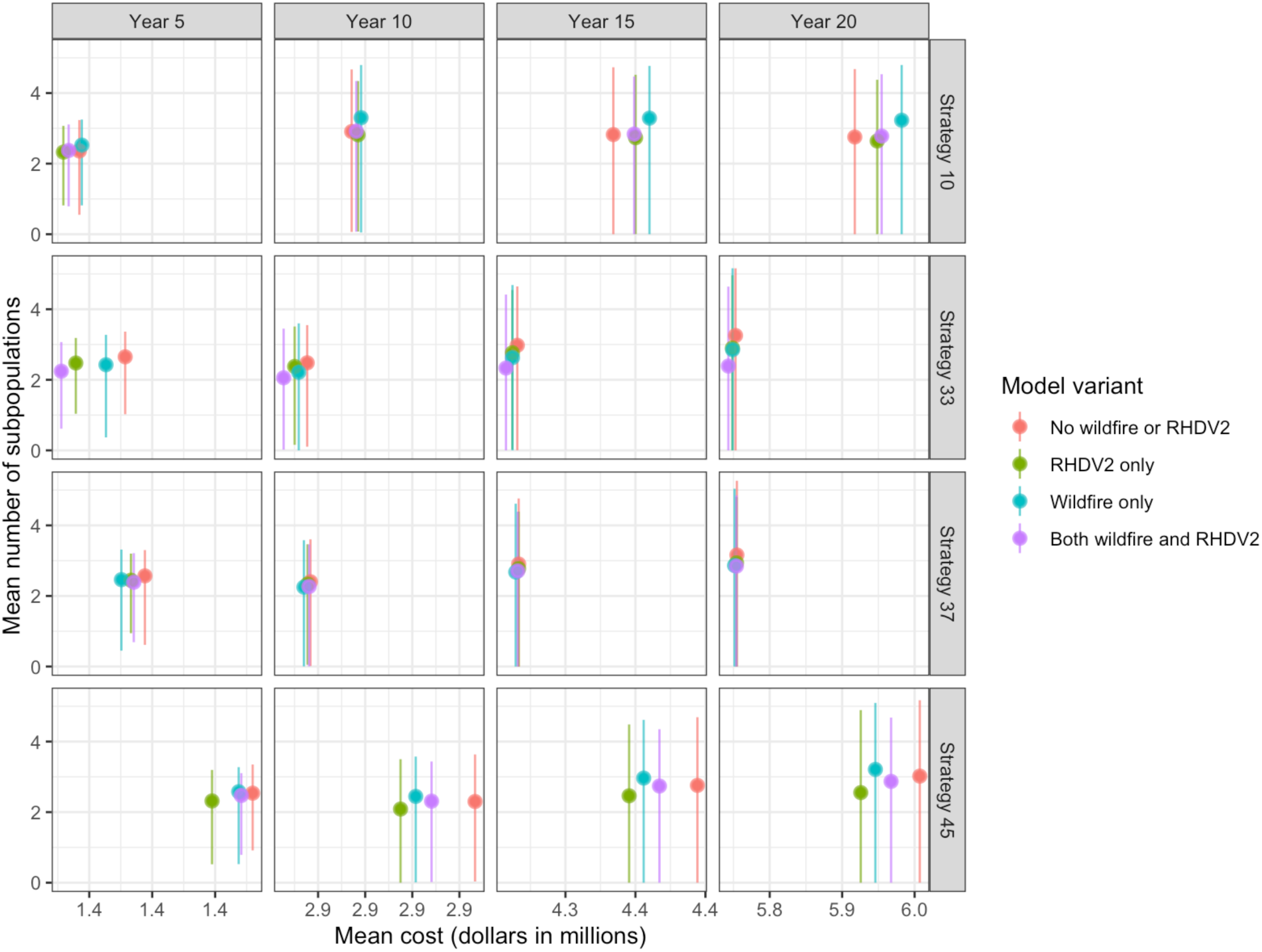
The horizontal panels display the mean number of wild subpopulations vs mean cost for each of the four strategies that repeatedly produced the highest mean number of wild subpopulations in multiple evaluation years, including the final year, under all variant conditions. The error bars indicate the upper and lower 95th quantiles for mean number of subpopulations across simulations in each evaluation year. Model variant is indicated by color, with two variants including RHDV2 outbreaks (green and purple), two including wildfires (blue and purple), and one excluding both types of events (red). Two strategies (the middle horizontal two panels above) performed best on average under the variant that excluded high impact events, while the other two strategies performed best in the variant that included only wildfires. All strategies had some risk of extinction by year 15 under all model variant conditions.

All strategies produced more wild subpopulations on average than the no intervention control strategy, which consisted of only the existing two subpopulations with no translocations between them, as well as no routine vaccination or breeding programs (identified in black in Figures 8, 9, 10, 11, and 12). The no intervention strategy had a lower cost than the strategies that incorporated at least one breeding program, but was comparable in cost with some strategies that did not have a breeding program.

**Figure 8.**
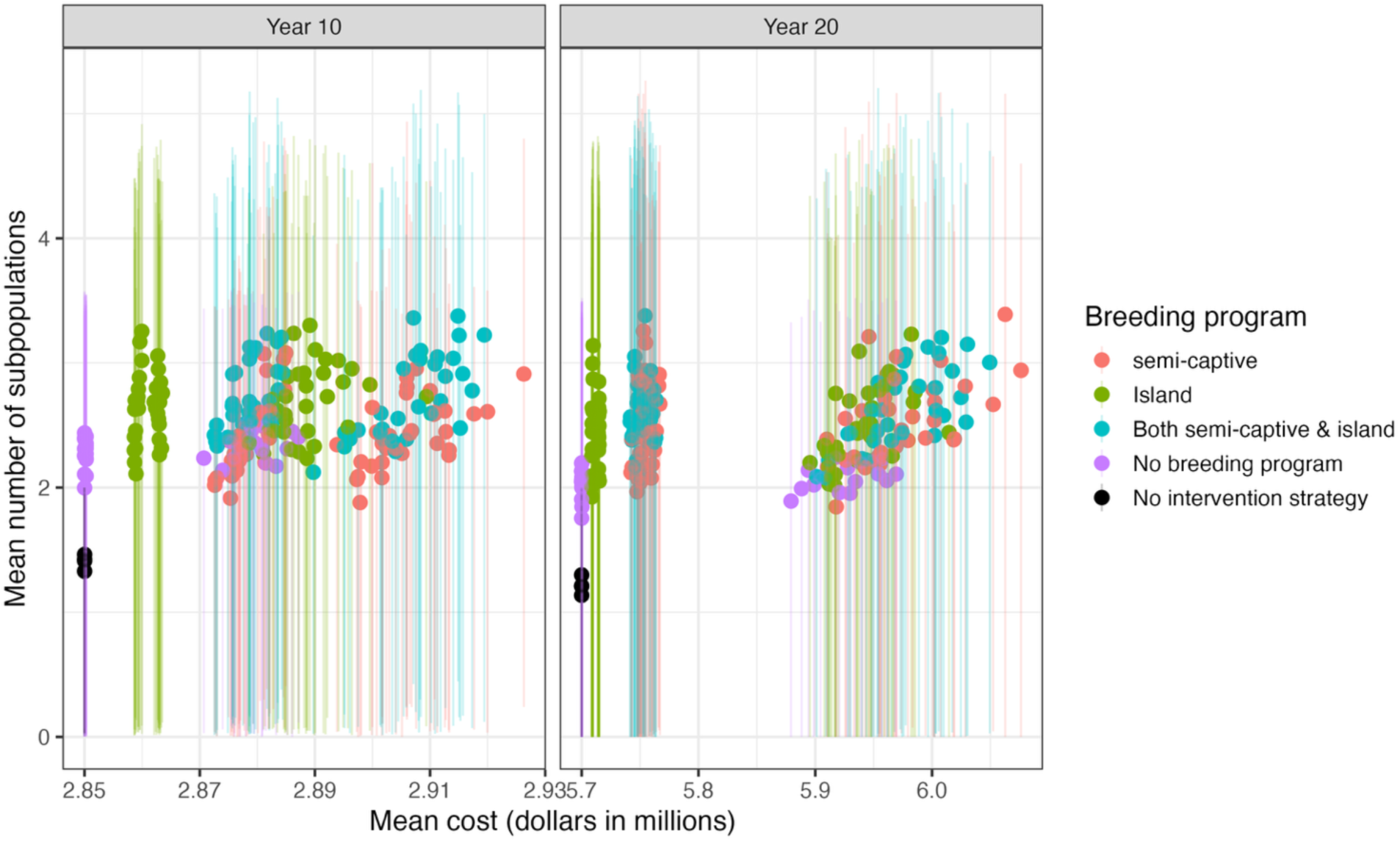
The mean number of wild subpopulations vs. mean cost of all strategies under all simulations, with error bars indicating the upper and lower 95th quantiles for mean number of subpopulations across simulations at the midpoint and final evaluation years. Many strategies have some risk of extinction by year 10, and all strategies have non-zero risk by year 20. A semi-captive breeding program is included in two action choices (red and blue), an island breeding program is included in two action choices (green and blue), and one action choice as well as the no intervention strategy have no breeding programs (purple and black).

Mean cost varied very little across strategies and through time, with the greatest difference between the lowest and highest mean costs occurring in year 20 ($264,258, approximately 4.6% of the lowest mean cost). The highest costs were due to vaccination of individuals at higher wild population levels.

#### 3.1.1 Conservation breeding program components

By year 10, the mean number of wild subpopulations resulting from strategies that incorporated at least one conservation breeding program was generally higher than the number resulting from strategies with no conservation breeding program, and this trend continued through year 20 (Figure 8). By comparing results for paired strategies in year 20, choosing to have at least one breeding program (semi-captive, island, or both) resulted in a mean of 0.49 more subpopulations (range-0.31, 1.29) and a mean of $37,967 higher cost (range: $59,351 lower to $158,194 higher) compared to having no breeding program.

Even in strategies with conservation breeding programs, however, the risk of extinction was > 0 by year 15. The choice of breeding program influenced cost, with higher costs for strategies that included a semi-captive breeding program in particular. By year 20, strategies with both breeding programs resulted in more subpopulations on average (0.11 more compared to semi-captive only [range-0.59, 0.69] and 0.17 more compared to island only [range-0.42, 0.79]). Strategies with both breeding programs were less expensive than those with only semi-captive programs (mean of $1,723 lower cost [range: $119,298 lower to $95,790 higher]) but were more expensive than island-only programs (mean of $36,855 higher cost [range: $70,979 lower to $114,994 higher]). By year 20, choosing a semi-captive breeding program resulted in a mean of 0.05 more subpopulations (range-0.84, 0.62) and a mean of $38,578 higher cost (range: $72,309 lower to $158,507 higher) compared to an island breeding program.

Only the semi-captive breeding program contributes directly to the cost calculation, through an annual probability of requiring enclosure fencing replacement due to stochastic events (wildfire, for example). Consequently, the difference in cost between the options that incorporate only semi-captive vs both semi-captive and island is not a significant difference, as it is driven by stochasticity in each simulation.

The choice of how frequently to retain juveniles in the breeding program to bolster the population had only a small effect on the number of wild subpopulations across all strategies (Figure 9). Strategies in which juveniles were retained every year generally had higher numbers of wild subpopulations in year 10, but by year 15 any discernible differences had disappeared. Options under this strategy component had no impact on mean cost. Comparing paired strategies in year 20 showed that choosing to retain juveniles every third year in one or both breeding programs resulted in a mean of 0.08 more subpopulations (range:-0.78, 0.83) and a mean of $3,027 lower cost (range: $106,397 lower to $116,694 higher) compared to retaining juveniles every year in one or both breeding programs.

**Figure 9.**
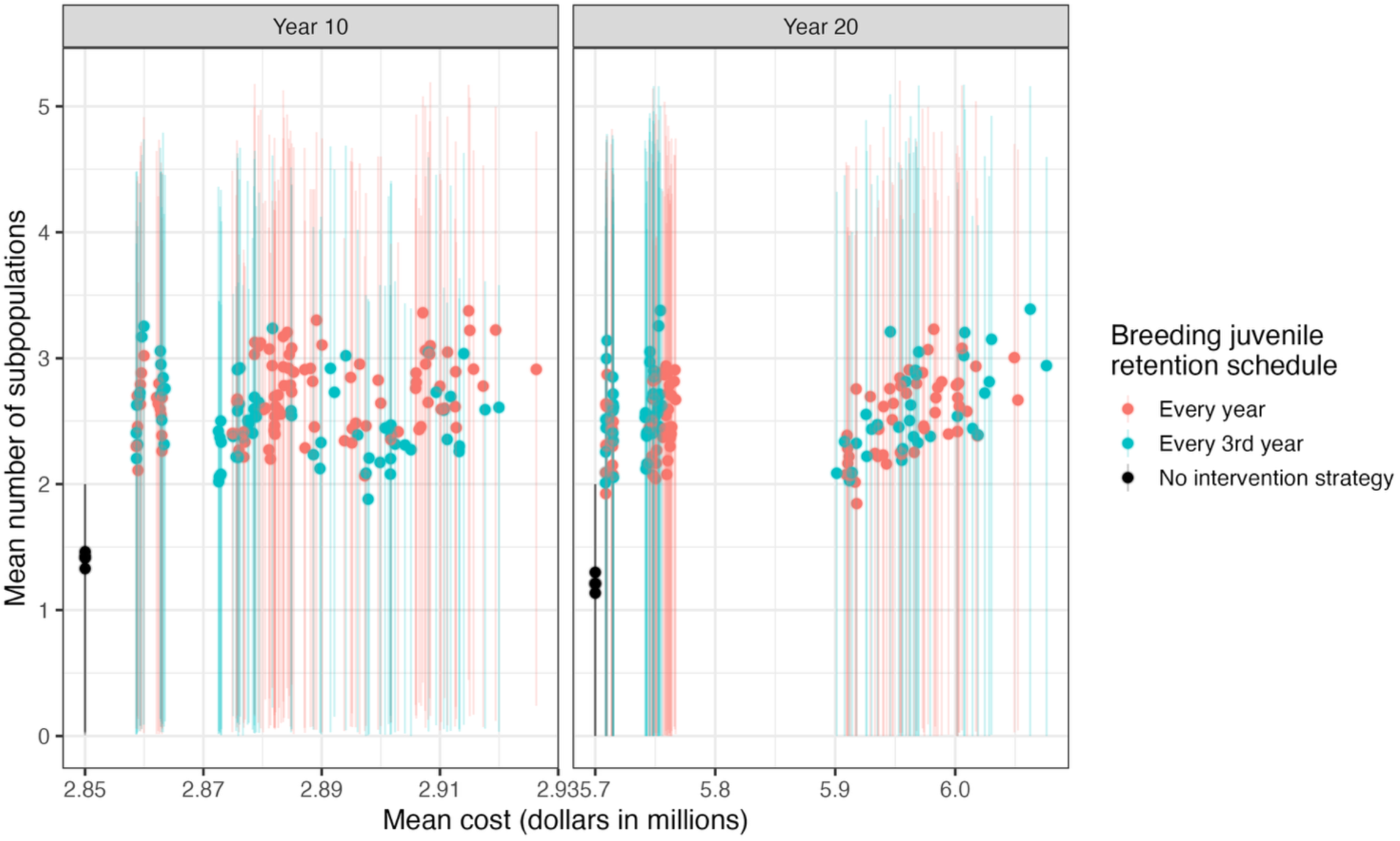
The mean number of wild subpopulations vs mean cost of all strategies under all simulations, with error bars indicating the upper and lower 95th quantiles for mean number of subpopulations across simulations for each strategy at the midpoint and final evaluation years. Strategies that retained juveniles in the conservation breeding program to bolster the breeding program every year are shown in red, and strategies that retained juveniles only every third year are shown in blue. Only strategies that included at least 1 breeding program were included in evaluating this action (48 strategies).

#### 3.1.3 Translocation component

The translocation component of how frequently to translocate individuals (every year or every 3^rd^ year) was only relevant in deciding between strategies when there were multiple types of wild subpopulations eligible to receive translocations, specifically subpopulations that were in the establishing phase (less than 5 years of occupation) and in crisis (fewer than five females left in the subpopulation in year *t*), and not enough available juveniles from the breeding program or wild subpopulations to translocate to all eligible subpopulations. The options – prioritize establishing or prioritize in-crisis sites – had no impact on mean cost and no consistent effect on the mean number of wild subpopulations through time (Figure 10). Prioritization of establishing sites resulted in the highest number of subpopulations in years 15 and 20, but the effect was relatively small and somewhat inconsistent across all strategies.

**Figure 10.**
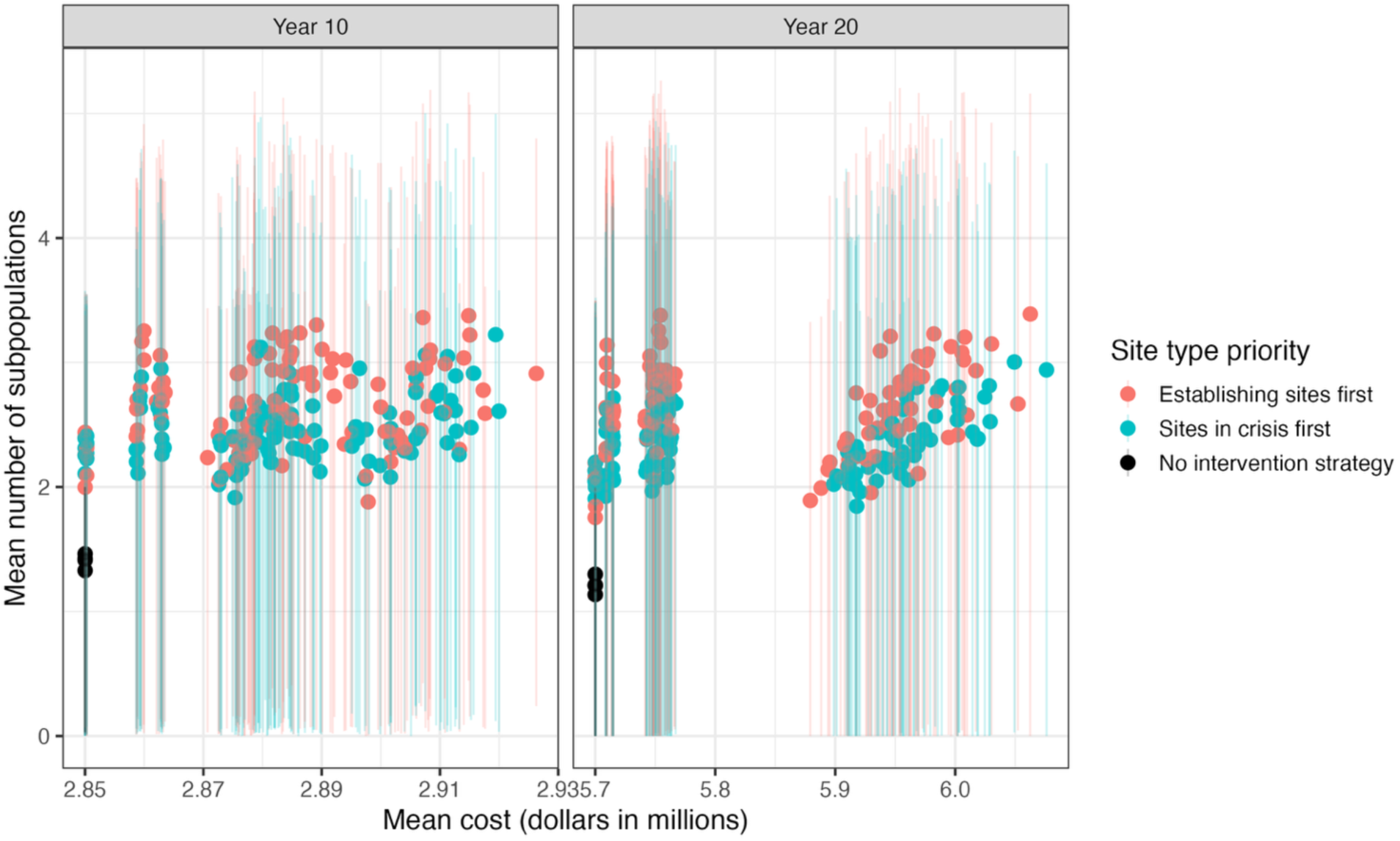
The mean number of wild subpopulations vs mean cost of all strategies under all simulations, with error bars indicating the upper and lower 95th quantiles for mean number of subpopulations across simulations for each strategy at the midpoint and final evaluation years. Strategies that prioritize translocations to establishing sites are shown in blue, and strategies that prioritize translocations to sites in crisis are shown in red.

Comparing paired strategies in year 20 showed that choosing to prioritize establishing sites resulted in a mean of 0.27 more subpopulations (range-0.35 to 1.20) and a mean of $1,386 lower cost (range: $101,112 lower to $107,112 higher) compared to prioritizing sites in crisis. Cost is not driven in any direct way by this component, however, and therefore the cost difference is likely driven by other components.

#### 3.1.2 Vaccination components

Vaccination options were evaluated only for model variant conditions that included RHDV2 outbreaks. These components only affected whether vaccinations were routinely administered in the conservation breeding program(s) or wild subpopulations; emergency vaccination procedures triggered by detection of an outbreak could occur regardless of strategy.

Routine vaccination in the conservation breeding program(s) did not have a substantial impact on the number of wild subpopulations across strategies that included a breeding program (Figure 11).

**Figure 11.**
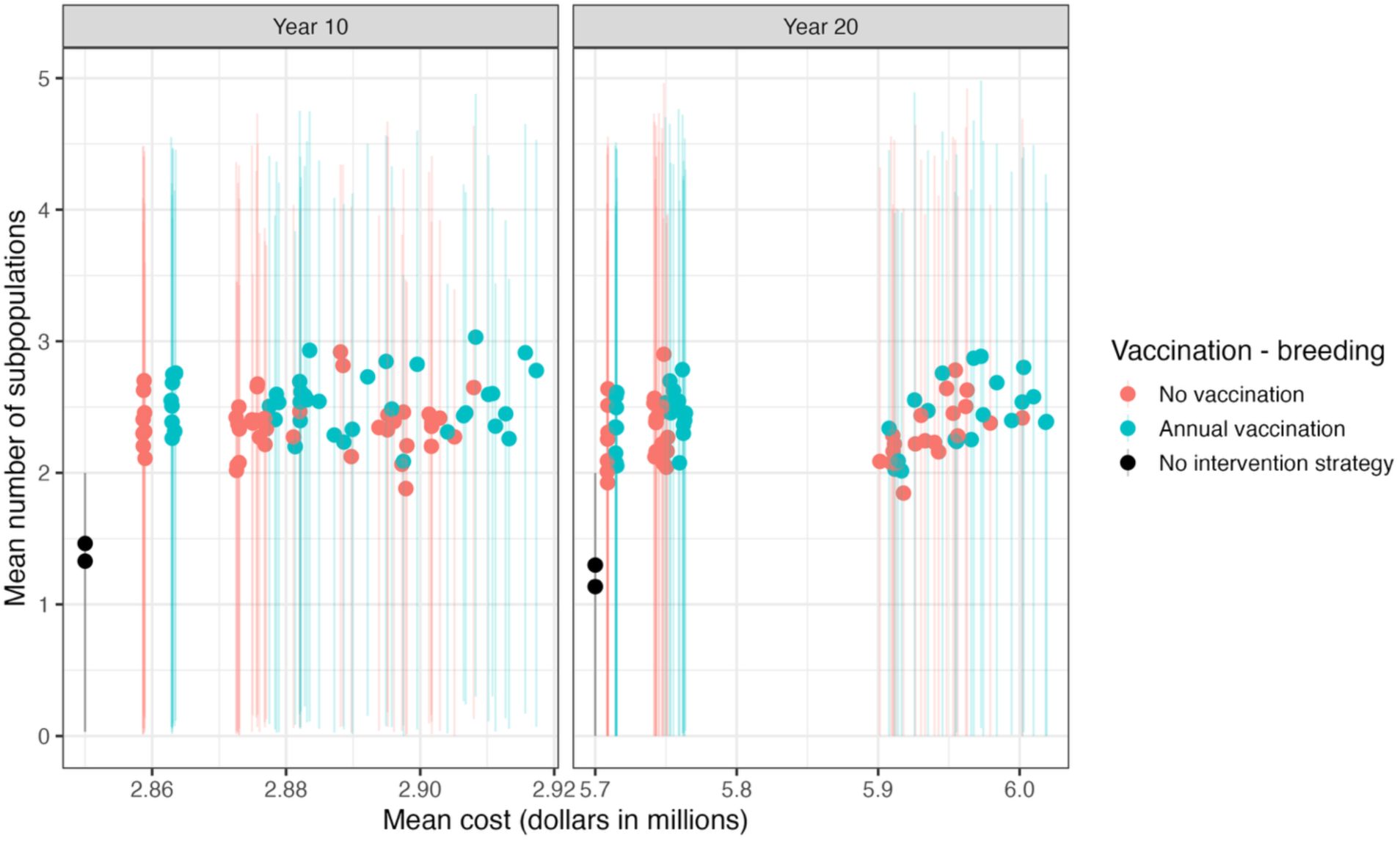
The mean number of wild subpopulations vs. mean cost of all strategies under all simulations, with error bars indicating the upper and lower 95th quantiles for mean number of subpopulations across simulations for each strategy at the midpoint and final evaluation years. Strategies that included routine annual vaccination of the conservation breeding program are shown in blue, and strategies that did not involve routine vaccination in the conservation breeding program are shown in red. Only strategies that included at least 1 breeding program were included in evaluating this action (48 strategies).

However, the highest costs in each evaluation year were associated with routine vaccination in the conservation breeding program(s). Comparing paired strategies in year 20 showed that choosing to routinely vaccinate in the conservation breeding programs resulted in a mean of 0.06 more subpopulations (range-0.54, 0.75) and a mean of $12,660 higher cost (range: $67,820 lower to $142,424 higher) compared to not vaccinating.

Routine vaccinations in the wild subpopulations also did not have a substantial impact on the number of wild subpopulations across all strategies (Figure 12). However, this strategy component was a primary driver of cost in year 10 and later. Comparing paired strategies in year 20 showed that choosing to routinely vaccinate in the wild resulted in a mean of 0.03 more subpopulations (range-0.39 to 0.54) and a mean of $226,437 higher cost (range: $148,912 to $318,219 higher) compared to not vaccinating in the wild.

**Figure 12.**
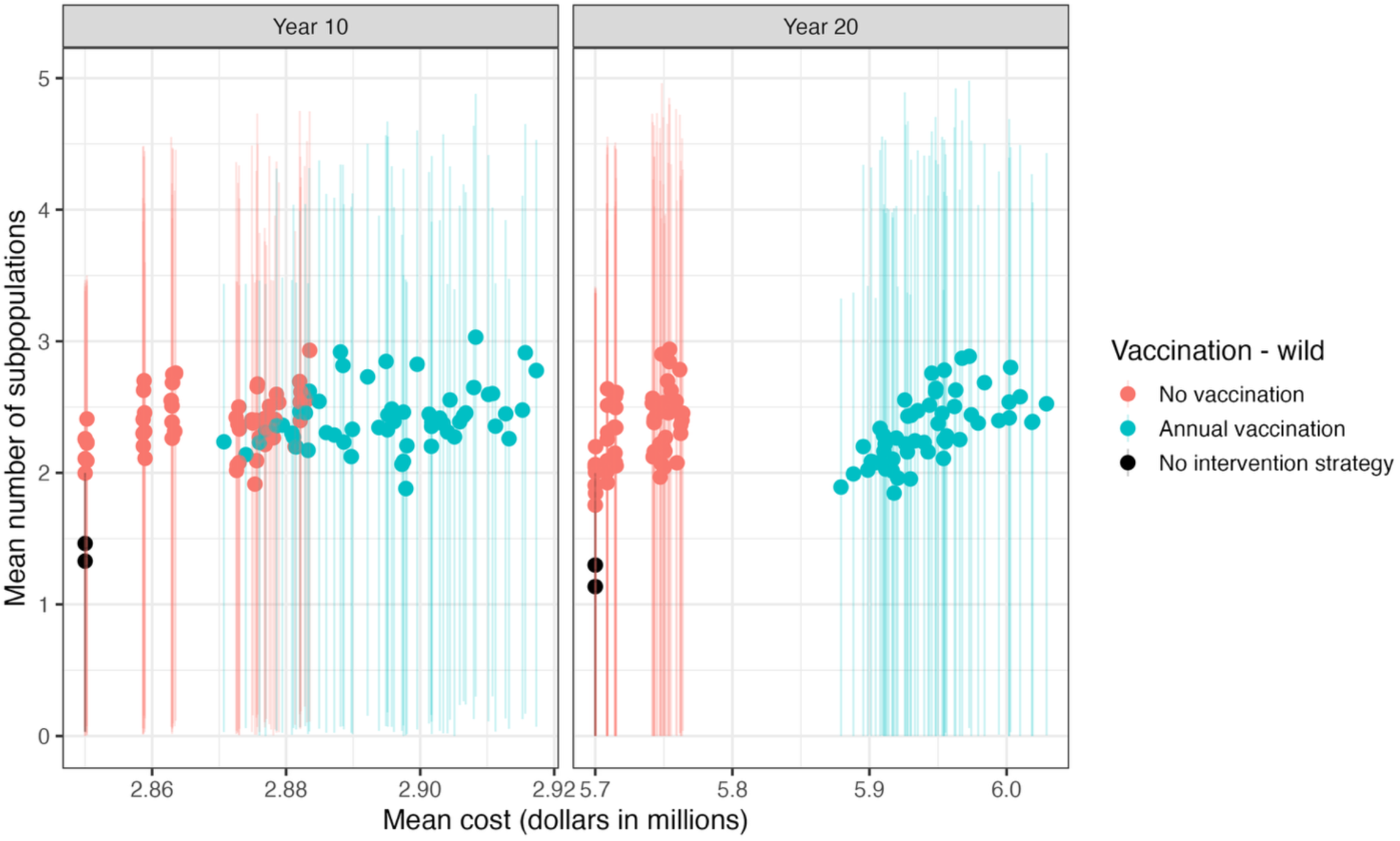
The mean number of wild subpopulations vs. mean cost of all strategies under all simulations, with error bars indicating the upper and lower 95th quantiles for mean number of subpopulations across simulations for each strategy at the midpoint and final evaluation years. Strategies that included routine annual vaccination of the wild population are shown in blue, and strategies that did not including routine vaccination in the wild population are shown in red.

#### 3.1.4 Impacts of wildfire & RHDV2 on population outcomes

For model runs that included RHDV2 outbreaks, the mean number of years with outbreaks across parametric simulations was 3.7. When sampling from the lowest 5% quantile of the annual outbreak probability, we observed a mean of 0.54 outbreak years out of 20 and when sampling from the highest 5% quantile, the mean number of outbreak years was 10.2. Both model variants that incorporated RHDV2 had lower mean numbers of wild subpopulations across all strategies, and the entire range of mean numbers of wild subpopulations per strategy was substantially lower than the variants that did not include RHDV2 (Figure 13). Mean extinction risk was also slightly higher for all strategies with model variants that included RHDV2, although the difference was not as visible for this metric because the range of mean values across strategies overlapped significantly for all model variants (Figure 14).

**Figure 13.**
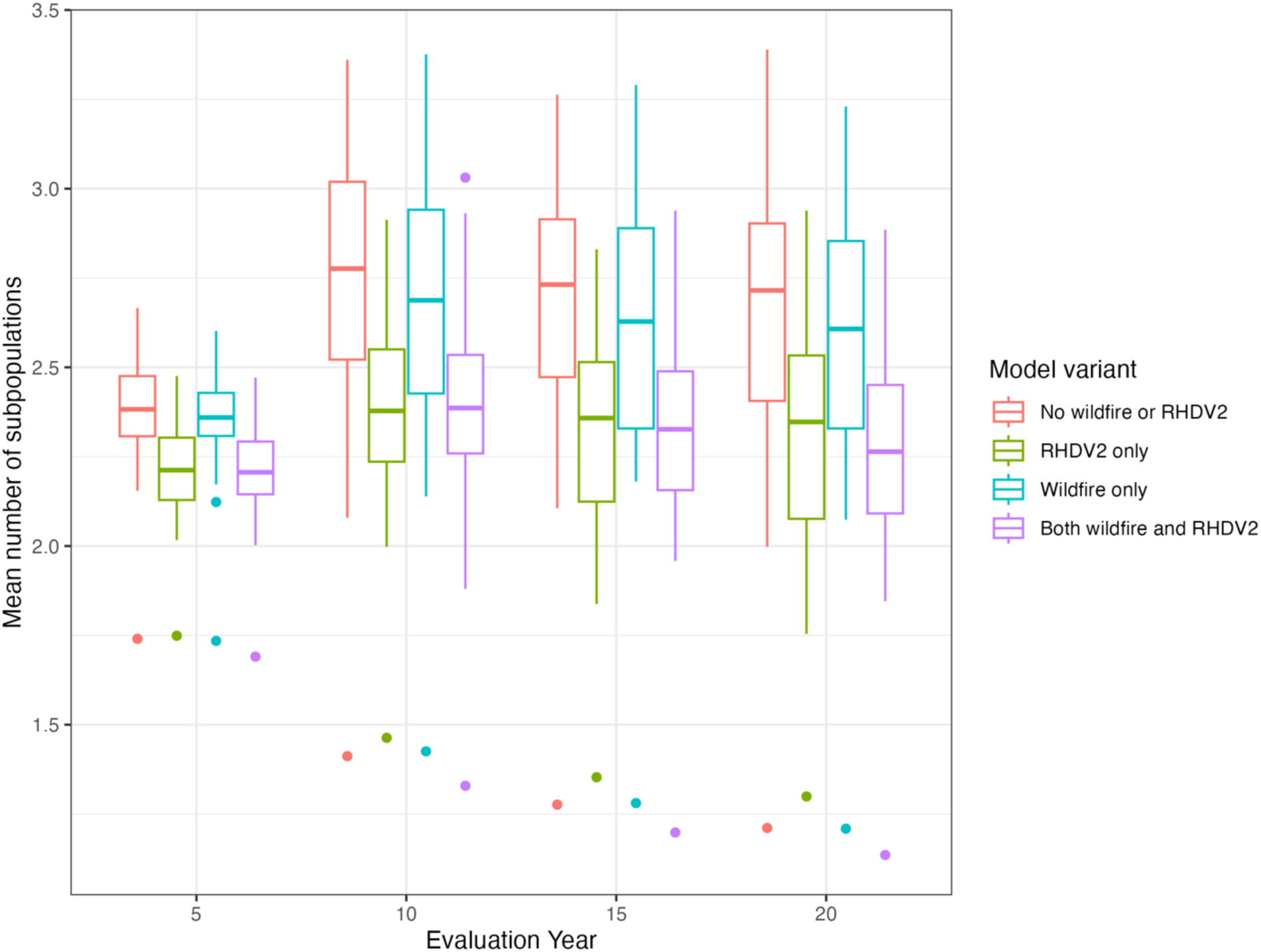
Boxplot of the range of mean numbers of wild subpopulations in each evaluation year produced by all strategies under each set of model variant conditions. Two variant settings include wildfires (blue and purple) and two include RHDV2 (green and purple) while one has neither type of event (red). The lower outlier points are the values from the no intervention strategy.

**Figure 14.**
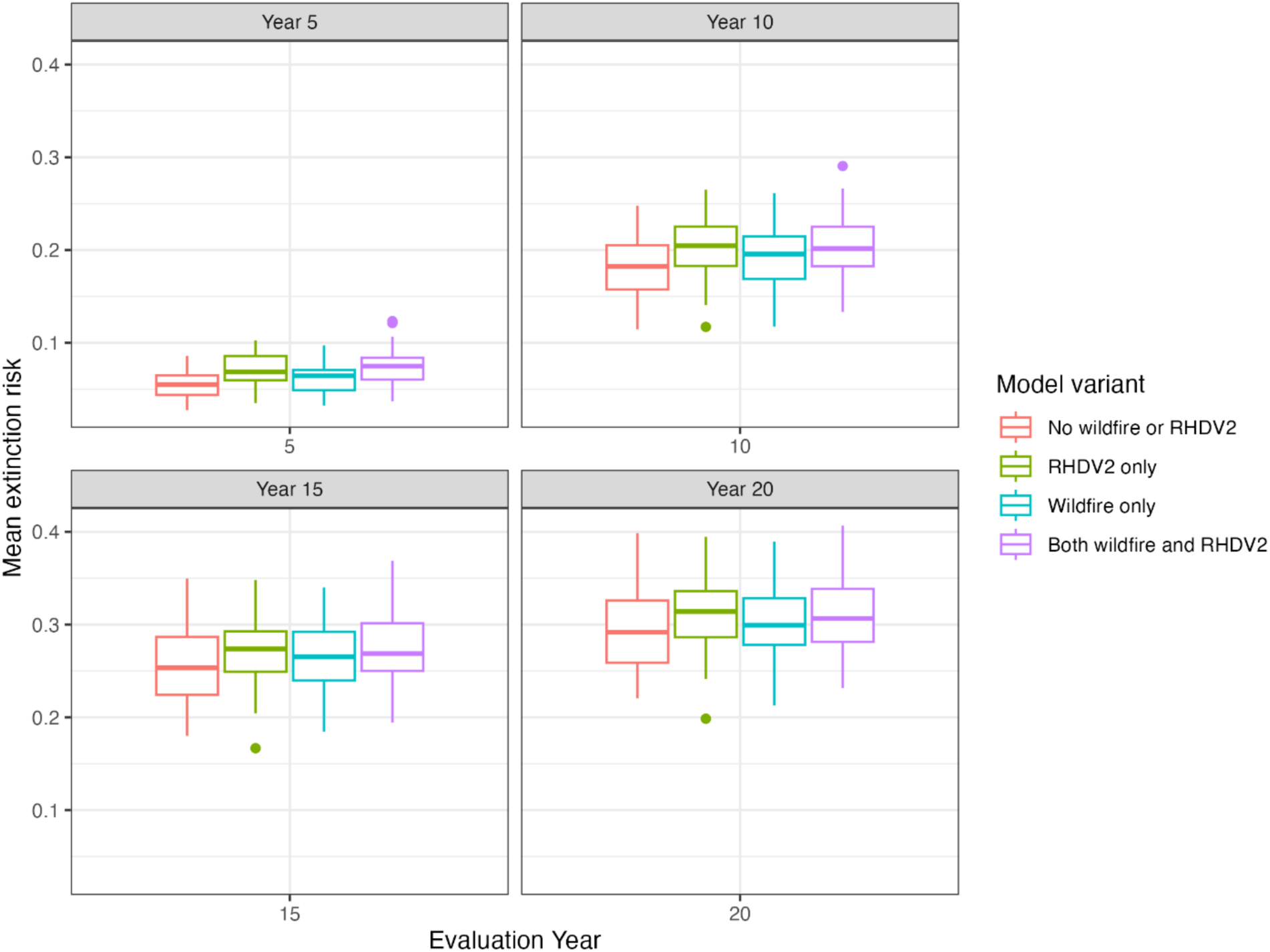
Boxplot of the range of mean extinction risk in each evaluation year calculated for strategies across all simulations, under each set of model variant conditions. Two variants included wildfires (blue and purple) and two included RHDV2 (green and purple) while one had neither (red). In years *t* = 5, 10, 15, and 20, extinction risk was calculated as the probability of extinction by year *t*.

Wildfire events had a much lower impact than RHDV2 outbreaks on the population outcomes under all strategies, both in terms of mean number of wild subpopulations (Figure 13) and, to a lesser extent, mean extinction risk (Figure 14). For model variants that included wildfire, on average there were 1.06 subpopulations affected by wildfires during the course of each 20-year model run. When comparing the distribution of the mean number of wild subpopulations produced by all strategies, there was substantial overlap between the model variant with only wildfire events and the model variant with no wildfire or RHDV2 outbreaks, although the overall mean value appeared to be slightly lower for the wildfire only variant, particularly in later evaluation years (Figure 13).

#### 3.1.5 Results comparison with the reduced wild survival scenario

Compared to the results presented above, the results from the reduced wild survival scenario showed, as expected, a decreasing mean number of wild subpopulations through time across all strategies (Figure 15) and higher extinction risks across time for all strategies (Figure 16) under all model variant conditions. Results from the reduced wild survival scenario had similar findings as above for four out of the five strategy components in terms of which option produced higher mean numbers of subpopulations. The only component that showed some difference in results was the juvenile retention schedule in conservation breeding programs. In the reduced wild survival scenario results, retaining juveniles every year was more strongly associated with the highest mean number of subpopulation values as well as with the highest cost values (Figure 17).

**Figure 15.**
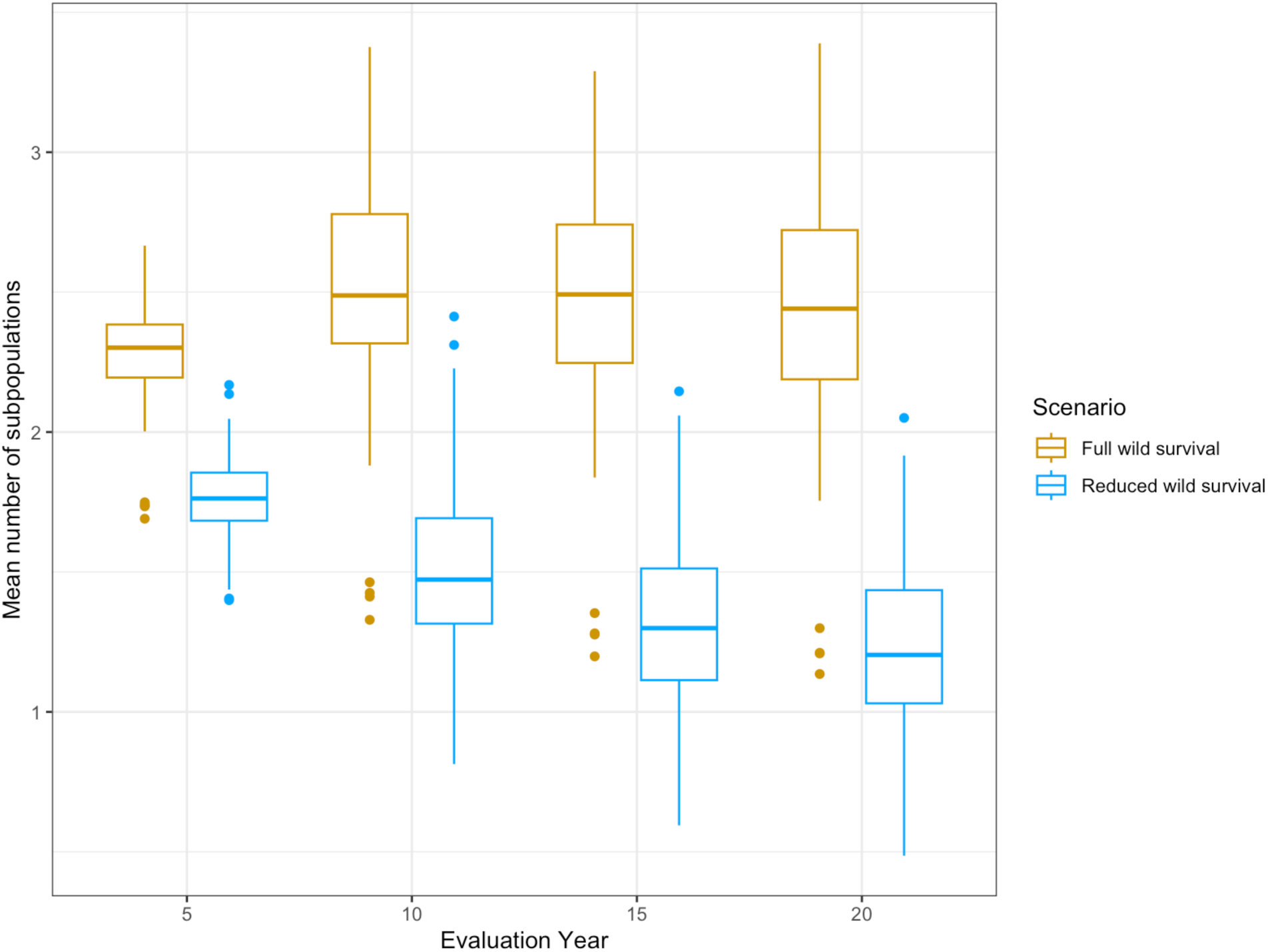
Boxplots of mean numbers of subpopulations across all simulations for all strategies under all four model variant conditions in each evaluation year for model results from the scenario with the full version of wild survival (gold) compared to the reduced wild survival scenario (blue). Reduced wild survival produced a generally declining mean number of subpopulations through time compared to the more stable mean through time of the full version.

**Figure 16.**
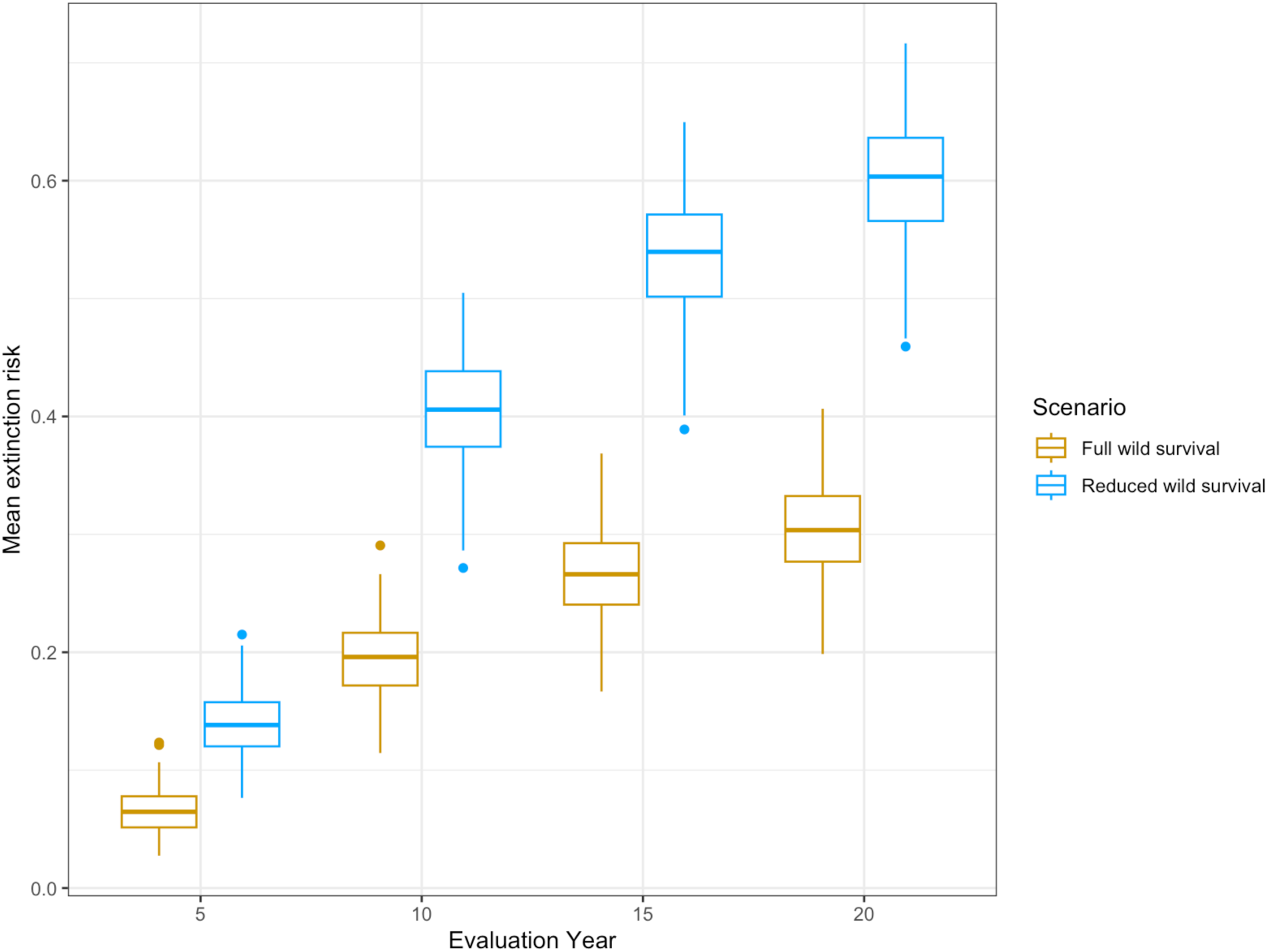
Boxplots of mean extinction risk across all simulations for all strategies under all model variant conditions in each evaluation year from the scenario with the full version of wild survival (gold) compared to the reduced wild survival scenario (blue). Extinction risk is cumulative in each evaluation year, and increases through time for both versions, but reaches much higher values by year 20 in the reduced wild survival scenario.

**Figure 17.**
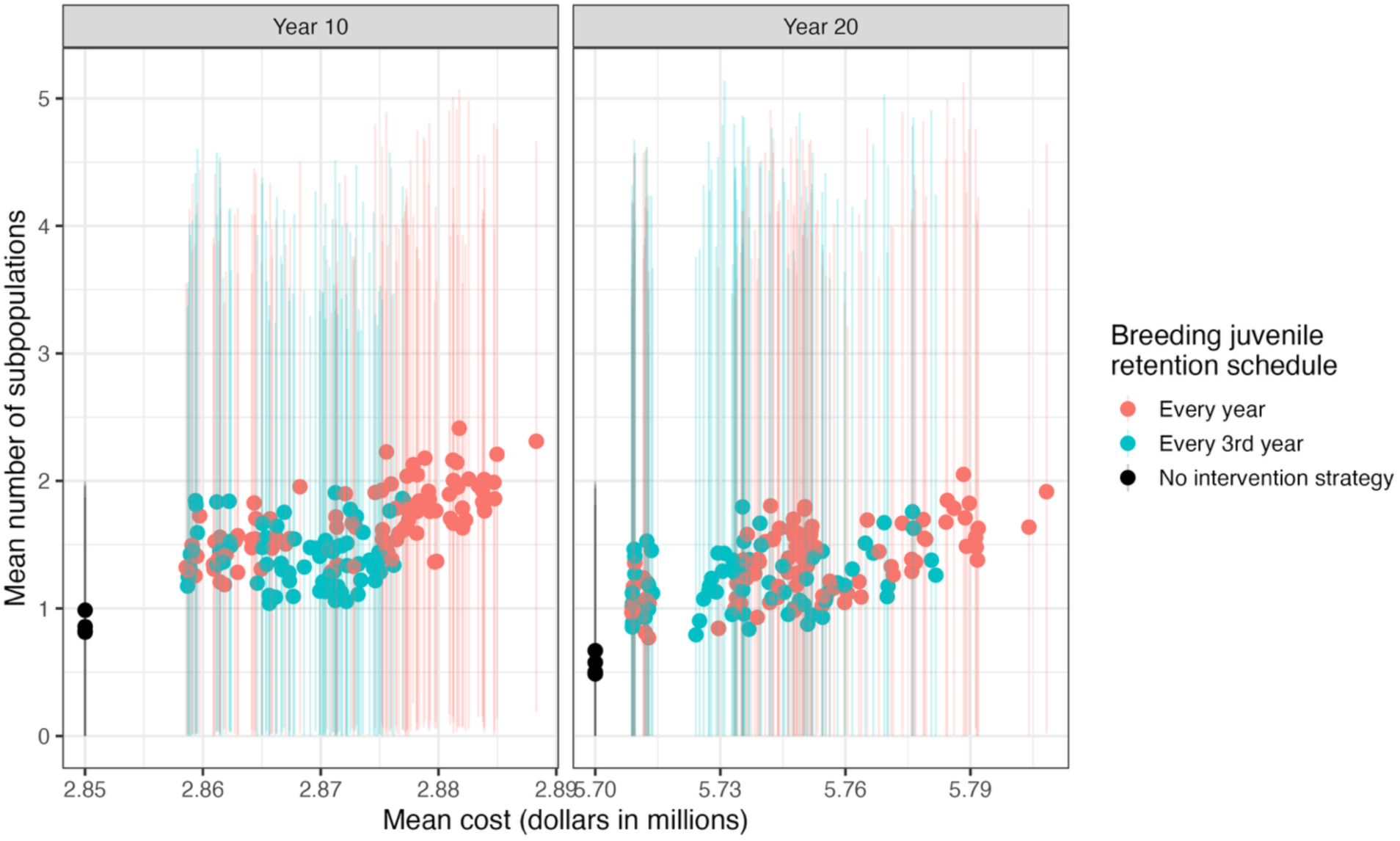
The mean number of wild subpopulations vs. mean cost of all strategies under all simulations, with error bars indicating the upper and lower 95th quantiles, at the midpoint and final evaluation years for the reduced wild survival scenario results. Strategies that retained juveniles in the conservation breeding program to bolster the breeding program every year are shown in red, and strategies that retained juveniles only every third year are shown in blue. Only strategies that included at least 1 breeding program were included in evaluating this action (48 strategies).

## 4 DISCUSSION

### 4.1 Key model assumptions and uncertainties

#### 4.1.1 Demographic estimates

The mean survival of adults was estimated at 0.34 (SE = 0.0002) in the wild and 0.33 (SE = 0.0001) in the conservation breeding program, which is in the range of what previous studies have found. DeMay et al. (2017) observed that 24 out of 113 (21%) CBPR adults survived to the first winter after being translocated to the wild from the semi-captive breeding program in the previous summer. Sanchez (2007) estimated female adult survival ranging between 0.09 (95% CI; 0, 0.23) and 0.44 (95% CI; 0.21, 0.67) across three wild subpopulations in Idaho. However, the fact that wild survival was estimated as higher than survival in the breeding program was a somewhat surprising result. Elias et al. (2013) estimated annual adult survival in the fully captive breeding program at 0.58 (SE = 0.04), higher than any wild estimate, although the conditions in the fully captive breeding program were different from conditions that occur in the semi-captive breeding program.

We evaluated how a version of this simulation model with lower wild survival, the reduced wild survival scenario, performed both in terms of overall population outcomes and objectives performance for each strategy component. We found that while there was certainly an impact on the simulated populations (in terms of both objectives, but particularly it produced a lower and declining mean number of wild subpopulations through time), there was less impact on the strategy component performance. Only one component, the breeding program juvenile retention schedule, showed some difference in relative performance of its two options under the reduced wild survival scenario (retention in every year was associated with the highest mean numbers of subpopulations) while with the version of the model using the full survival coefficient distributions, the two options (retention every year versus retention every third year) had very similar performance.

#### 4.1.2 Lack of natural migration

A key assumption in the model is that there is no significant natural migration between current wild subpopulations, and, given the locations of the current and proposed recovery area sites, it is unlikely to occur more frequently in the future. There have been a few instances of individuals moving between Beezley Hills and Sagebrush Flats in the past, but it is very likely that direct management actions will be required for the foreseeable future to establish new wild subpopulations and potentially to maintain the genetic diversity of all subpopulations. In the model, there is no mechanism for individuals to move between wild sites on their own; estimating movement probabilities between populations would likely be very challenging.

#### 4.1.3 RHDV2 and wildfire impacts

RHDV2 outbreaks and wildfires are events with high uncertainty and potentially very high impacts, and as such are key sources of uncertainty in the model outcomes. Several sources of uncertainty about RHDV2 outbreaks were incorporated in the model through parametric uncertainty (through the annual probability of outbreak and the probability of advanced detection in the event of an outbreak), and RHDV2 was shown to have more impact on the mean number of wild subpopulations compared to wildfires. However, parametric uncertainty about wildfire impact was not directly incorporated into the model (though stochasticity was).

While the probability distributions for the number and size of fires were estimated using data, fire size in particular was difficult to model because there were a couple of very large fires that constituted outliers, such that the data did not fit any standard statistical distribution well. Excluding these fires was not reasonable because wildfires in the future will likely be larger and higher intensity under climate change (Cunningham, 2024, Iglesias et al. 2022). Instead, we grouped fire size into size classes with homogeneous effects within each class. In order to avoid setting a specific upper limit for fire size, fire zones were created, and a fire of size class 3 was assumed to affect any subpopulation within the same fire zone. Additionally, any subpopulations affected by wildfire was assumed to have 100% mortality, which represents a conservative approach. In reality, fires that could potentially impact pygmy rabbit populations based on proximity may not actually reach them depending on the direction of the fire. Even if there is some impact of a fire, a fire may not affect the entire subpopulation or the intensity of the fire may allow for some survival even within the affected subpopulation. In sum, these model choices may have led to a positive bias in wildfire effects.

On the other hand, fire zones were based primarily on a single satellite map of the area and therefore represented a relatively coarse approach to understanding potential fire spread. In the model, fire zones served as hard barriers to fire spread, which prevented fires adjacent to an RA zone from affecting it if the fire originated in a different fire zone than the RA site, regardless of the size of the fire. It is not known whether this could have led to an underestimation of the effect of large fires. Additionally, probability distributions for both the number of wildfires per year and fire size class were static through the 20-year model runs and therefore do not reflect any increase in the frequency and intensity of wildfires relative to the available data. The data do indicate a trend towards more fires per year, and some indication of increasing frequency of larger fires, between 2008 and 2024. These model choices may have led to a negative bias in wildfire effects.

### 4.2 Evaluation of management strategies

Based on the performance of the alternative strategies in terms of the number of wild subpopulations, results indicate that continuing to operate a breeding program will increase the likelihood of expanding into and maintaining new wild subpopulations. The model results indicate that the semi-captive breeding program is particularly important, likely because it is already established, but that having both a semi-captive and island breeding program would be beneficial. Since this result occurred even when wild survival estimates were on average higher than the breeding program, it is likely that simply having the redundancy of a conservation breeding program increases the ability of the species to survive stochastic catastrophic events, such as wildfires and RHDV2.

In the reduced wild survival scenario results, the positive impact of conservation breeding programs on the number of subpopulations was even more apparent. These results reinforce the importance of a breeding program for a scenario in which wild survival is generally lower than breeding program survival, with the same emphasis on the importance of the semi-captive breeding program in particular. Based on the results from this scenario, there may also be a small advantage associated with retaining juveniles to support the breeding program every year, rather than every third year.

#### Model results also support continuing to administer annual RHDV2 vaccinations

While the amount of impact on the mean numbers of wild subpopulations was not enormous or uniform across strategies, the highest mean numbers of subpopulations under model variant conditions that included RHDV2 were primarily associated with strategies where annual vaccination occurred in the breeding program, and comparing paired strategies showed a small increase in the number of subpopulations on average. The wild vaccination component did not have an impact on the number of wild subpopulations, but this seems likely to be due to the relatively low proportion of the population that was subject to vaccination in any given year (30%). This proportion was set based on the low estimate from WDFW managers for the current proportion of the wild population that they are able to vaccinate in any given year (30 – 50%). Further experimentation with this model could be done by manipulating that proportion to determine if there is a tipping point at which the vaccinated proportion of the wild population is high enough to make a substantial difference in the number of subpopulations.

The highest mean cost values were associated with annual vaccinations, with the largest differences associated with vaccinations in the wild population. The relative difference in cost between strategies was very low, however, only slightly more than 1% of the total at the greatest point of different (∼$67,000 out of $5.7 million in year 20).

Routine vaccination impact in the models may have had less apparent impact on the model results because all strategies (including the no intervention control strategy) had emergency vaccination automatically occurring if an RHDV2 outbreak was detected. While we incorporated multiple sources of uncertainty into that process, an important constraint that was not included was resource limitation. In reality, an emergency vaccination process will be constrained to some extent by the available resources, in terms of money and personnel, and this may not match our model assumptions for the potential effectiveness of an emergency response to RHDV2. In which case, routine vaccinations will provide the primary and perhaps only protection against RHDV2.

Model results provide moderate support, in terms of the number of wild subpopulations, for prioritizing the translocation of juveniles into new recovery areas during their establishing phase (the first five years) over wild subpopulations that have declined to a very low level (less than 10 individuals).

While this strategy component was not the only contributing factor, the strategies that achieved the highest number of subpopulations in years 10, 15 and 20 were those that prioritized establishing sites. Comparison of paired strategies also showed an increase in the mean number of wild subpopulations by year 20 by prioritizing establishing sites, with no increase in mean cost.

Finally, our results support revising the design of the monitoring program to more effectively estimate demographic rates in both the wild subpopulations and in the conservation breeding program, including annual survival and reproduction, and annual variation in these vital rates. Understanding how survival and reproduction vary by age class, wild versus breeding program, as a function of habitat, and over time, is critical for making more informed management decisions. The reduced wild survival scenario showed that the results of one management action was affected by a small adjustment to wild survival (constraining the parameter distribution to ensure that wild survival was lower than breeding survival), and illustrates the importance of continuing to improve our understanding of demographic rates. Our results here are conditional on the demographic information that was available, and that information could be made more robust with changes to how the population is monitored. All information on reproduction came from a couple of years of monitoring (2012 – 2014) in the semi-captive population.

Furthermore, the current monitoring design does not facilitate estimating survival or how it varies with space, time, or other factors.

### 4.3 Other management opportunities

There were many more possible management actions that were brainstormed as part of the SDM process that were not suitable to directly incorporate into the population model. These actions included ideas aimed at increasing the amount of suitable habitat available to pygmy rabbits, increasing protections for existing and potential recovery areas, and increasing support for pygmy rabbit conservation through community engagement and conservation partnerships.

Suitable habitat for pygmy rabbits has both biotic and abiotic requirements. While the abiotic requirements (primarily soil composition suitable for digging burrows) are difficult to affect through direct management, biotic requirements may be more easily manipulated. Increasing the amount of suitable habitat in existing and future recovery area sites could be accomplished with management actions such as:

- Habitat enhancement in proposed recovery area sites by augmenting perennial grasses, forbs and shrubs using native (local ecotypes) seed sources and controlling invasive weeds
- Undertaking habitat restoration projects in burned portions of existing recovery areas (Mansfield Plains)

One challenge that was identified by the group regarding increasing suitable habitat for pygmy rabbits is the fact that, while we can reliably identify some characteristics of suitable habitat (soil composition and density and sagebrush stand maturity and density in particular; WDFW 1995, USFWS 2007), in past translocations of CBPR, individuals abandoned sites that were identified as suitable by managers and thrived in areas that were considered marginal (WDFW biologist J. Gallie, personal comm.). Given this inconsistency, further study into the finer details of what characterizes suitable habitat for pygmy rabbits is warranted to identify what other actions might lead to habitat enhancement.

Existing and proposed recovery areas are threatened by natural processes such as wildfires as well as human development. There are actions that can be taken to lessen the impact of wildfires on CBPR populations, including:

- Increasing wildfire protection in rabbit areas through direct habitat manipulation (e.g. fire barriers around occupied habitat)
- Protecting unprotected lands by incorporating them into fire districts
- Including rabbits as “values at risk” for wildfire response Protections against human development can be pursued in the form of:
- Enhancing CBPR knowledge to inform land-use authority of local jurisdictions (e.g., via the WDFW Priority Habitat and Species program)
- Promoting Conservation Reserve Program retainment to ensure lands already in that program remain
- Incorporating CBPR considerations into Programmatic Biological Opinions (e.g., renewable energy)

Engaged communities and conservation partners are key to the long-term support of conservation initiatives, and continually developing these relationships is an important aspect of conservation. For CBPR conservation, this is particularly crucial because some of the proposed recovery areas are partly or entirely on private lands and will require landowner cooperation to develop as recovery areas. In order to accomplish this, enrolling new landowners in Conservation Benefit Agreements (CBAs) and renewing existing CBAs may be required. In addition, other actions to engage local communities in pygmy rabbit conservation could include:

- Advertising volunteer survey opportunities to local community members
- Regularly updating the community on CBPR conservation news and developments (print and broadcast news, social media, etc.)

Actions to develop new or enhance existing relationships with conservation partners could include:

- Expanding coordination through the CBPR working group
- Connecting with the Washington Shrubsteppe Restoration and Resiliency Initiative (WSRRI) to use this as a forum for collaborating on habitat management actions for CBPR
- Expanding and strengthening coordination with Tribes, conservation districts (CDs), and non-governmental organizations (NGOs)
- Delivering shared messages through networks (e.g., Tribes, CDs, agencies, private lands biologists)
- Maintaining internal communication and support for the CBPR program within WDFW and USFWS

### 4.4 Future model extensions and possible modifications

Further research into CBPR survival and reproduction would be beneficial, particularly the difference in reproduction and survival between wild and semi-captive individuals, and variation in reproduction and survival over time, to continue to evaluate the value of the breeding program into the future. Furthermore, a better understanding of juvenile survival, which could not be reliably estimated with the current data, is critical to understanding population growth.

Future study on how translocations affect survival and reproduction (*sensu* Armstrong et al. 2017) would also be valuable, as this could influence future management. In the model currently, there is no survival or reproduction cost associated with any type of translocations. While this choice was made based on expert judgment, it is an assumption that merits further study to identify and mitigate any significant costs to survival, given that translocations are essential to establishing new wild subpopulations and to maintaining the genetic diversity of the breeding program and all wild subpopulations.

In terms of model choices, one that merits future reconsideration is the creation of fire zones to limit the spread of very large fires. One possible alternative approach that could be experimented with to better incorporate large fires would be to attempt to fit a more flexible distribution to the size of fires in the WDNR dataset. Another alternative would be to use the fire class sizes as they are currently defined and set an upper limit for fire size in size class three.

## APPENDIX A: SDM PROCESS RESULTS

### Decision framing statement

The USFWS and WDFW are continuing to work to maximize the viability and progress toward recovery of the federally and state-listed Columbia Basin Pygmy Rabbit (CBPR). Much of the management of the CBPR in recent decades has been through management of and releases from the semi-captive breeding program, in order to re-establish and increase the population size in the wild, resulting in two surviving wild subpopulations (subpopulations as defined in USFWS SSA). Fostering landowner support through Conservation Benefit Agreements (CBAs) has been prioritized and was vital to this re-establishment, and expanding the population into new areas will require collaborating with new landowners. Other types of actions have been taken, such as habitat management and improvement, vaccination, and others. However, the effectiveness of these actions for the wild population has been challenging to evaluate. The semi-captive populations have struggled to produce in recent years and have themselves declined significantly, providing an opportunity to re-evaluate the program’s objectives and methods.

Threats to the rabbit both in the wild and in the semi-captive program include low population size, effects from climate change, the associated risk of catastrophic events such as wildfires, habitat loss from wildfires, agricultural conversion and development, and disease. The decision makers would like to develop an adaptable strategy, which is sustainable and implementable for guiding management in the coming decades across the range of the CBPR, taking into account changing conditions and updated information. Primary challenges associated with this decision include ecological uncertainty, limited resources for management, and the constraints of working within the mosaic of primarily private lands in the recovery area.

### Objectives

Fundamental objectives were identified by the group in three categories; objectives related to directly measuring the performance of the pygmy rabbit population, objectives related to conditions that would support the pygmy rabbit population (habitat and public support), and objectives related to the cost associated with supporting pygmy rabbit management.

- Objectives directly measuring the pygmy rabbit population:

`o` Maximize the persistence (resilience) of CBPR
`o` Maximize redundancy (number of subpopulations) across the historical range of the CBPR
`o` Maximize the ecological diversity of the CBPR (representation of multiple types of suitable habitat - higher elevation)
`o` Maintain the genetic signature of the CBPR (representation)
- Objectives for conditions supporting the pygmy rabbit population:

`o` Maximize high-quality CBPR shrubsteppe habitat within CBPR range
`o` Maximize landowner support for CBPR
`o` Maximize general public support for CBPR

- Objectives related to the cost of pygmy rabbit population management:

`o` Minimize the level of management intensity for CBPR
`o` Minimize cost and human resources devoted to CBPR

In addition to the above fundamental objectives, the group identified strategic and means objectives for the pygmy rabbit management program:

- Strategic objectives

- Establish and maintain healthy, resilient, robust, self-sustaining CBPR populations in WA
- Improve sustainability of conservation program & activities
- Means objectives:

`o` Increase the number of wild populations across a larger portion of the shrub steppe landscape (decreasing risk of one fire taking them all)
`o` Improve understanding of disease risks (RHDV, tularemia, plague, unknowns) for informed management decisions (vaccines, flea control, etc)
`o` Evaluate the feasibility of, and as appropriate, implement habitat improvement to bolster wild rabbit populations
`o` Identify and secure habitat in conservation status
`o` Focus habitat restoration/improvement funding/actions in areas (private/public) we proactively want to move rabbits in the coming decade.
`o` Improve understanding of CBPR habitat requirements & tolerances
`o` Initiate outreach to new landowners to increase conservation benefit agreement (CBA, formally SHA) enrollment to expand private lands areas for pygmy rabbit reintroductions
`o` Maximize the reproductive capacity of the conservation breeding program
`o` Optimize monitoring protocol to assess wild population abundance & spatial distribution
`o` Minimize disruption of agricultural activities and land use
`o` Maximize public/private landowner support of rabbit recovery
`o` Minimize public/private landowner concern of ESA regulations
`o` Foster support for pygmy rabbit conservation efforts from county commissioners & nearby communities
`o` Minimize disease transmission & risk to CBPR through education to larger community
`o` Maximize research to address information gaps in CBPR outside of just genetics - applied questions are particularly important, like how they are dispersing, moving, responding to environmental factors
`o` Increase confidence in our decision-making, so that “scrambling” doesn’t occur
`o` Develop adaptive management processes
`o` Inform and update the WA state recovery plan

### Alternative strategies development

Many strategy components were considered during the course of the SDM process. Some components were converted into model choices after receiving feedback from SDM participants (1b, 2b, 2c, 4a and 4c), while one was incorporated into the model as a source of uncertainty (1a) after preliminary results did not show that the options of that component had an impact on the objectives model outcomes.

#### All strategy components considered (final components used in strategies bolded)

1. Developing new RA sites

a. Rate of site development annually

i. Low, medium or high (1, 2 and 3 sites per year respectively)
b. Site selection criteria (based on site evaluations in Gallie et al. 2024)

i. Logistically simplest, least amount of habitat restoration required, spatially distributed throughout the historical range of CBPR
2. Breeding program

a. Which breeding programs to include

i. **Semi-captive only, island only, both or none**
b. Initial action in year 1 for juveniles currently in the semi-captive breeding program

i. Remain in semi-captive breeding program or release into a wild subpopulation
c. How many subpopulations to maintain in the semi-captive breeding program

i. 1 to 4
d. How frequently to retain juveniles in breeding program(s) for maintenance

i. **Every year or every third year**
3. Vaccination

a. **Wild population**

i. **Annual vaccination or none**
b. **Breeding program**

i. **Annual vaccination or none**
4. Translocations

a. Type of site priority

i. Establishing only, establishing and in crisis, or establishing, in crisis and declining
b. Preferred site if more than one type of site can be a priority in the event there are not enough juveniles to go to all priority sites

i. **Establishing first or in crisis first**
c. Distribution strategy for translocating juveniles to more than 1 site

i. Equally between eligible sites or proportional based on the area of the sites

## APPENDIX B: EXPERT ELICITATION RELATED TO RHDV2

Rabbit hemorrhagic disease virus (RHDV2) is an emerging disease that has caused very high mortality in both wild and domestic rabbit populations across the western US in the last five years. While there have been some reported cases of outbreaks in Washington state in 2024 (in San Juan County), no outbreaks have yet occurred that could affect the pygmy rabbit population, and therefore few data were available to construct probabilities of an outbreak occurring or the likely impact an outbreak would have on the CBPR. The decision-making group decided that explicitly including the risk and potential impact of RHDV2 outbreaks in the model was critical for evaluating management options, particularly whether to vaccinate against RHDV. An expert elicitation process was undertaken to decide how to best incorporate the risk and impact into the SDM model and construct the necessary probability distributions.

The experts consisted of a subset of the decision-making group, including members representing both USFWS and WDFW, as well as Dr. Janet Rachlow, a professor at the University of Idaho and member of the CBPR Science Advisory Group. During initial discussions with the group, it was decided that any RDHV2 outbreak that occurs in the pygmy rabbit population will likely result in the exposure of the entire population, both in the wild and in any breeding programs that may exist, and therefore all individuals will be subject to disease-related mortality (with the specific risk dependent on vaccination status) if an outbreak occurs. The primary sources of uncertainty that the elicitation focused on were the annual probability of an outbreak occurring, whether there would be enough time to mount an emergency vaccination response and for emergency vaccines to become fully effective, and the added annual mortality that the wild population might experience as a result of captures related to vaccination.

The expert elicitation process followed a modified Delphi method (MacMillan and Marshall 2006), consisting of two rounds of responses to questions with a meeting to discuss results between the rounds. Each round consisted of each expert responding individually to questions in an online form. The final versions of the questions were:

- What is the annual probability of an RHDV2 outbreak occurring within the pygmy rabbit population in the historical range of CBPR?
- What is the probability of detecting an RHDV2 outbreak if it is occurring within a 100-mile radius of the historical range of CBPR?
- What is the possible absolute increase in annual mortality from captures related to vaccination? Each expert was asked to give their best, minimum and maximum probability values for each question (between 0 and 1), along with their confidence (50 to 100%) level that the true value occurred within the range they gave for each round (Spiers-Bridge et al. 2010).

Because all of these parameter estimates were probabilities, i.e., bounded by 0 and 1, we fit beta distributions to the resulting data. The parameters of beta distributions, for each question and expert, were estimated by minimizing the sum of squares of each expert’s quantiles and best values. All distributions were then standardized to a 99% confidence interval and combined to create a mixture distribution for each parameter (as described in Sipe 2023) which was then used in the population model.

## Results

The combined distribution of all experts’ responses for the annual probability of RHDV2 outbreak had a mean value of 0.19 with a long right tail to a maximum value of 0.95 (95th quantiles were 0.03 and 0.56). Responses varied substantially between experts, indicating a high degree of uncertainty about this parameter (Figure B1).

The combined distribution for the probability of detecting an RHDV2 outbreak before it appears in the pygmy rabbit population indicated even higher uncertainty, with a mean value of 0.49 and a range of 0 to 1 (95th quantiles: 0.04, 0.95). Some experts estimated an essentially uniform distribution for this probability, while one estimated a fairly low probability and two others estimated fairly high probabilities, producing a bimodal mixture distribution (Figure B2).

When estimating how much added annual mortality may occur from capture myopathy due to vaccination operations in the wild population, two experts estimated much higher rates of mortality than the other four (up to 0.45), which produced a mixture distribution with values that, when added to the mean baseline annual mortality produced by the survival analysis (0.76 for wild), could produce an annual survival of 0 and cause the population to collapse. To avoid this, we decided to remove the two highest experts’ distributions from the combined distribution. The combined distribution with the remaining four experts’ responses is bimodal, with one mode centered around 0.0001 and the other around 0.001 (Figure B3).

**Figure B1.**
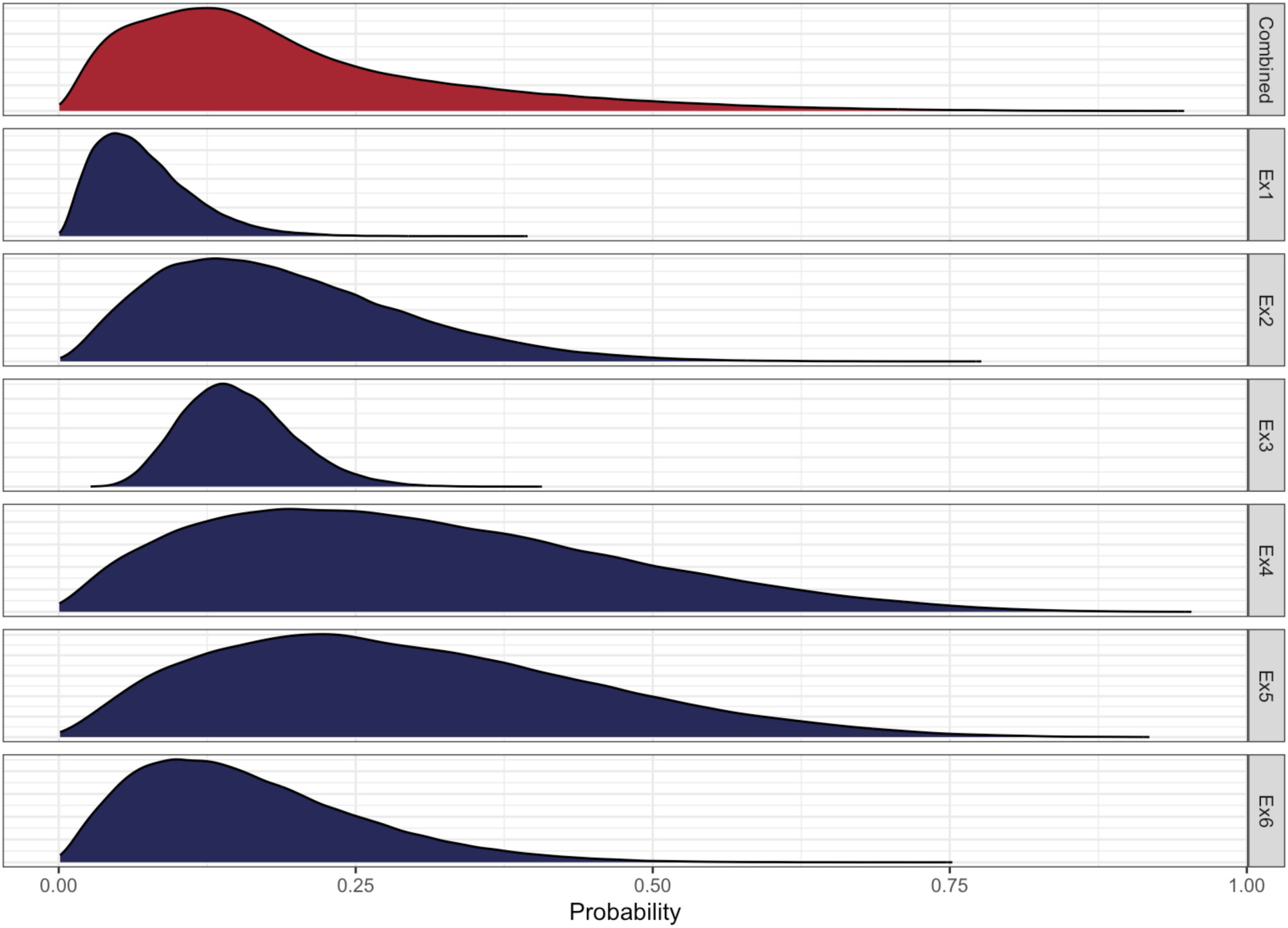
The standardized distributions for the annual probability of an RHDV2 outbreak of each of the six experts (in blue) and the resulting combined distribution (in red).

**Figure B2.**
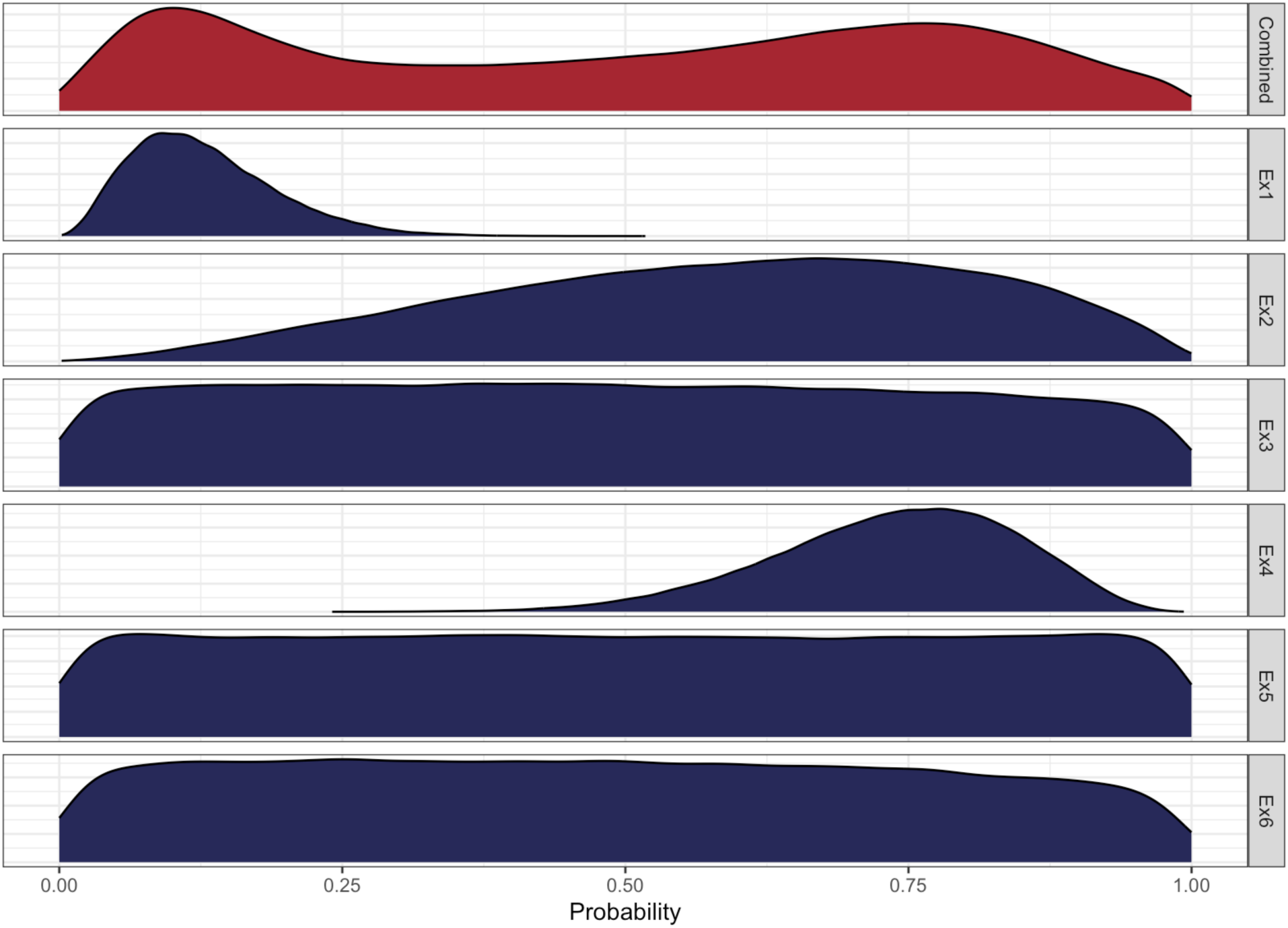
The standardized distributions for the probability of detecting an RHDV2 outbreak before it appears in the pygmy rabbit population (given that an outbreak occurs) of each of the six experts (in blue) and the resulting combined distribution (in red).

**Figure B3.**
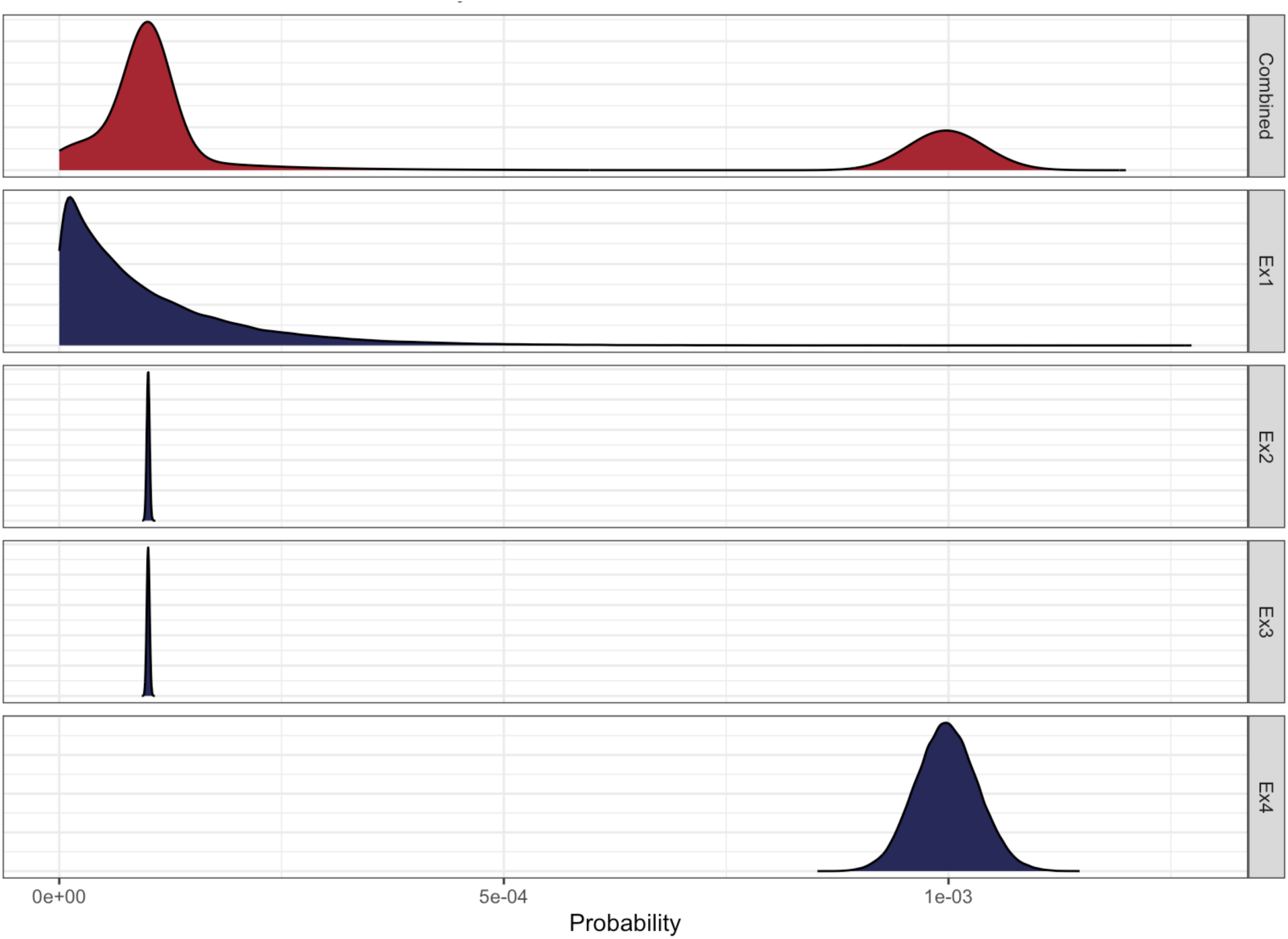
The standardized distributions for the added annual mortality associated with vaccinations (due to capture myopathy) in the wild population of each of the six experts (in blue) and the resulting combined distribution (in red).

## APPENDIX C: SURVIVAL ANALYSIS

### Data description

Monitoring data were collected between 2012 and 2024 in both the semi-captive breeding program and the wild subpopulations. For analysis, the dataset was subset from 2016 to 2024 because the methods used in the semi-captive breeding program changed from permanent enclosures to semi-captive enclosures in 2017, which were found to be a better method for preventing disease outbreaks in the conservation breeding program. Individuals were uniquely identified through DNA, collected either during live captures occurring in either location type, or using pellet sign sampling during winter surveys conducted in wild locations (see Figure C1).

The age of individuals was either determined through live capture observations and extrapolated from there based on the first observation date or extrapolated based on the time between observation dates. In the model context, a year begins at the start of the breeding season on March 1, with the first litters of the year typically appearing beginning in April. For individuals labeled as juveniles based on a live capture observation, any observations up to the March following that observation were labeled juvenile, and all observations after that March were labeled adult. For individuals with no observed age but with observations made more than 2 years apart, the first observation date was labeled juvenile, and any observations after the March following that date were labeled adult (only 5 individuals met this criteria). For individuals that were observed multiple times but less than 2 years apart, any observations after the March following the first observation were labeled adult, and observations prior to that were labeled unknown age and excluded.

The dataset was then subset to individuals that had at least 2 observations (removing 577 out of 806 individuals), since including individuals with only one capture event made it difficult for the model to separate detection probability from survival probability. However, we note that this has the potential to result in a positive bias in survival. After removing 20 more individuals with unknown ages or unresolvable date entry errors, there were 229 individuals included in the analysis observed over 311 observation days between 1/1/2016 and 10/10/2024. Observation days did not occur at the same time of year through time for live trapping, which is the source of the bulk of the observations (Figure C2), while pellet sign surveys occurred in the winter, but only in the wild.

Individual capture histories were constructed using the location of each individual through time, matched to the observation dates of either observation method that occurred in that location. For individuals that were translocated, their last possible observation date in the origin location was used as the translocation date, and starting on the day after the translocation date they were considered available for detection in the new location. A cap on the last possible observation date was set to 5 years after the first observation of the individual (the maximum recorded age for a pygmy rabbit is 5.7 years; Elias et al. 2013), to shorten model runs while avoiding impacting survival estimates. Out of the 311 total observation days in the dataset, the number of observation days that any given individual was available for detection, based on the location of the individual and the locations where observation effort occurred on each day, ranged between 3 and 73 observation days.

**Figure C1.**
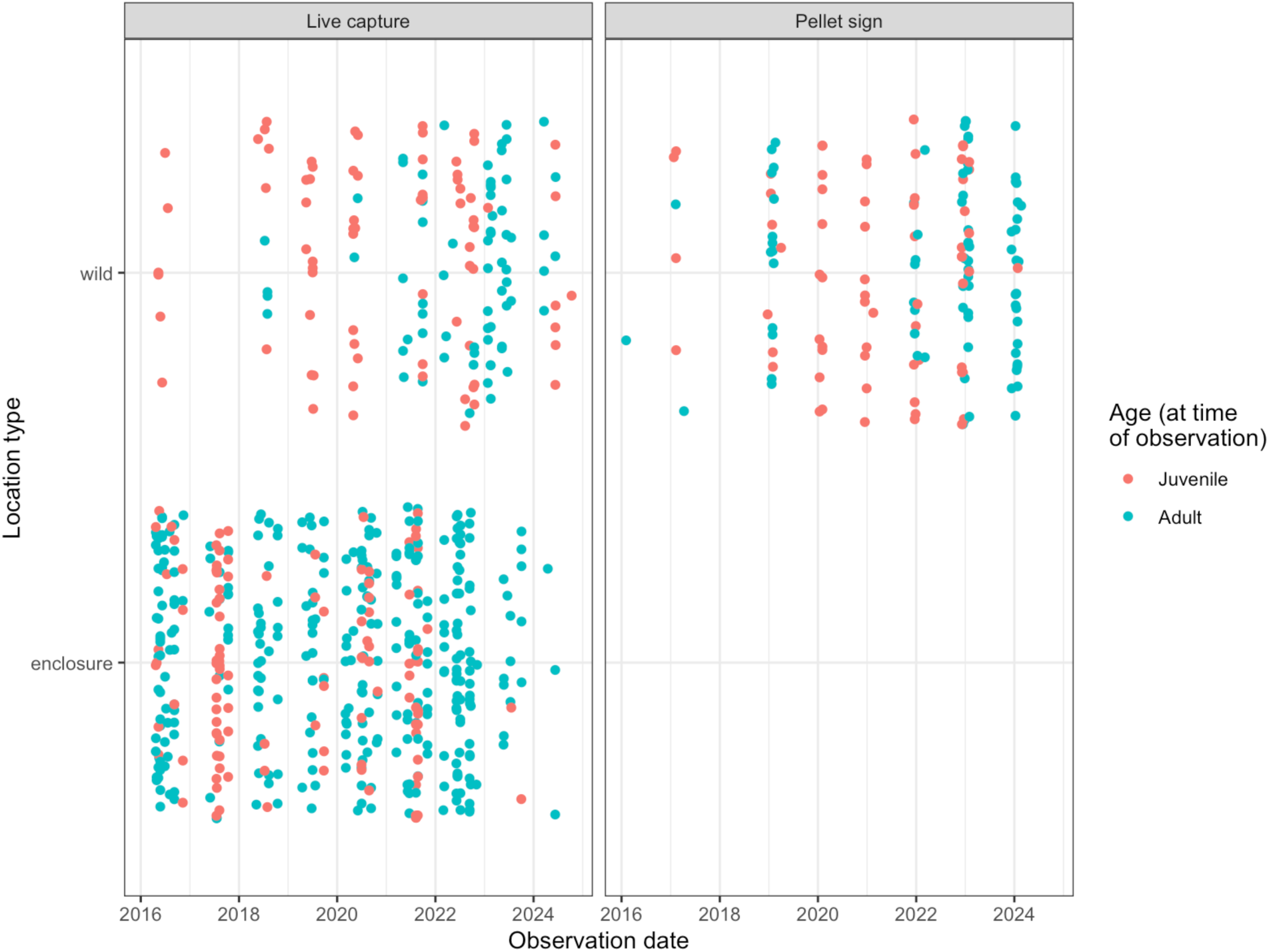
Observations of individuals through time by observation location type (wild or semi-captive enclosures) for each detection type (facets). Age at the time of observation is indicated by color, red for juveniles and blue for adults. Many of the observation days shown above for live capture of juveniles in the wild were translocation releases and did not count as observation days in that location (instead it was an observation day in the origin location, where individuals were trapped in order to translocate them).

**Figure C2.**
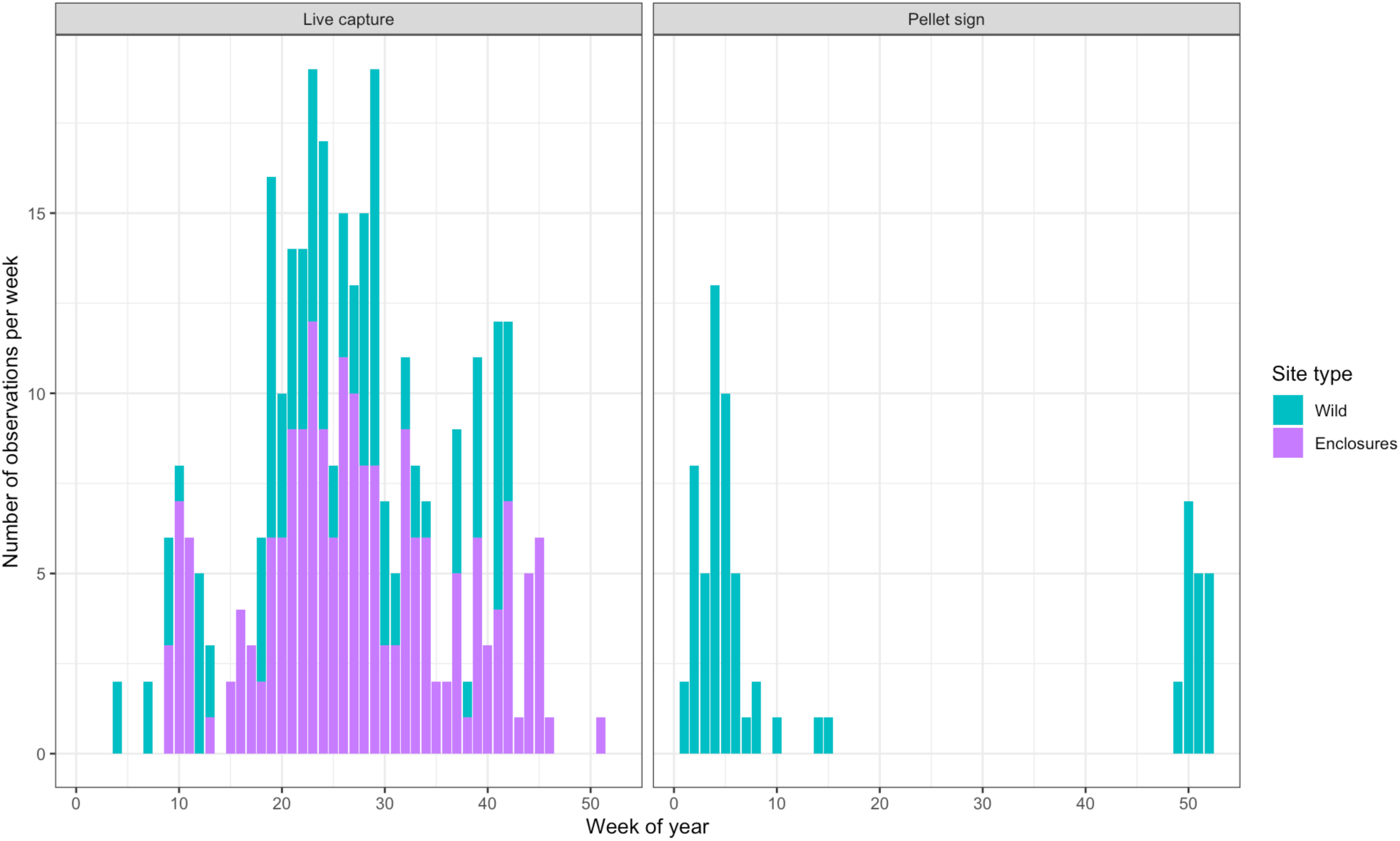
The number of observations in each week of the year across all years for live capture (left plot) and pellet sign sampling (right plot) between 2016 and 2024, in wild locations (green) and in the semi-captive breeding program (purple). Pellet sign is clustered in the winter and could potentially have been used alone to estimate annual survival directly, but this method was only used in the wild. Live trapping was used in both locations but does not cluster temporally through the years in a way that would make estimating annual survival directly easy.

### Survival model

A capture mark-recapture model was used to estimate survival and detection probabilities.

Survival probability is modeled as function of age (juvenile or adult) and location type (wild or conservation breeding program) while detection probability is modeled as a function of detection type (live capture or pellet collection) and location type.

The true state of an individual (*z* = 1, alive, or *z* = 0, dead) on sampling day 𝑗 + 1 is modeled as a Bernoulli outcome with an estimated daily survival probability for individual 𝑖 from observation day 𝑗 to 𝑗 + 1 (𝜙_*,)_), raised to the power of the number of days (𝑑) between day 𝑗 and 𝑗 + 1:

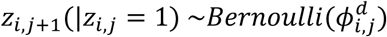

Daily survival probability (𝜙*_i,j_*) is then modeled with a logit link and a linear model including an intercept (𝛼_𝜙_), fixed effects of age (𝑎), and location type (𝑙), and a random effect of observation day (𝜃):

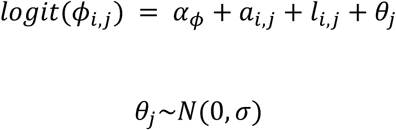

The prior distribution of the standard deviation of the random effect is Uniform(0,1), and the priors for the intercept and coefficients for wild location type and juvenile age are Normal(0,1). Adult age and breeding location type coefficients were set to 0. The intercept therefore describes the adult breeding survival as the baseline.

Detection of individual *i* on sampling day 𝑗 + 1 is modeled as a Bernoulli outcome with a daily detection probability (𝑝_*,)_):

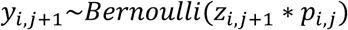

Daily detection probability (𝑝_*,)_) is modeled with a logit link and a linear model including an intercept (𝛼_7_), and fixed effects of detection type (𝑑) and location type (𝑙):

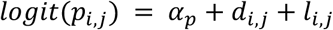

The prior probability distributions for the intercept and fixed effect coefficients for wild location type and live capture detection type are Normal(0,1). Breeding location type and pellet sign detection type coefficients were set to 0. The intercept therefore describes the breeding location pellet sign detection as the baseline.

The survival analysis was conducted using the Nimble package (version 1.2.1), and model diagnostics were performed using the MCMCvis (version 0.16.3) and coda (version 0.19-4.1) packages in R.

## Results

All parameters for both survival and detection probability converged well (Figure C2 and C3), with 𝑅^S^ values < 1.05 (calculated based on Gelman and Rubin 1992). Distributions of survival probability were generated by drawing 10,000 samples from the intercept, fixed effect coefficients, and the standard deviation of the random effect, then survival was calculated for each combination 20 times with random draws from the corresponding random effect distribution, for a total of 200,000 survival probabilities

The model estimated juvenile survival as much higher than adult survival; the mean of the distributions are 0.67 for juveniles in both location types (SE = 0.0003 for wild, 0.0002 for breeding enclosures), and half that for adults, 0.34 (SE = 0.0002) and 0.32 (SE = 0.0001) for wild and breeding enclosures respectively (see Table C1 for all coefficients mean and SE estimates). The wild location type coefficient is more widely distributed than the breeding enclosure type coefficient for both age groups (Figure C4).

## Discussion

The survival probability estimates produced for adults in this analysis are similar to estimates from other survival studies of pygmy rabbits; DeMay et al. 2017 observed 21% apparent survival of adults in the wild after translocation in one year and Sanchez (2007) estimated female adult survival between 0.09 (95% CI; 0, 0.23) and 0.44 (95% CI; 0.21, 0.67) for three different wild subpopulations in Idaho. However, juvenile survival estimates were extremely high compared to observed survival from other studies; DeMay et al. 2017 reported that the proportion of juveniles translocated to the wild from the semi-captive breeding program in the summer that survived to the following winter was 0.39, 0.13 and 0.10 in 2012, 2013 and 2014 respectively. There were many juveniles in the group of individuals that were excluded from the analysis because there was only 1 observation, many of which were translocated from the breeding program to the wild (the translocation event was the only observation). It seems likely that this exclusion meant that survival was biased high for juveniles, since juveniles that were observed at least twice were the ones that survived long enough for this to occur. Due to this bias, the juvenile survival estimates from the analysis were not included in the population model, and instead juvenile survival was set to 75% of adult survival in each location type in each year in the model, to reflect the lower survival that juveniles typically experience compared to adults.

The average adult survival in the wild was estimated by the model as being slightly higher than adult survival in the conservation breeding program, which was a surprising result. The variance in wild adult survival was also higher than for breeding program adults, however, and this combined with the fact that there were relatively few wild adult observations in the analysis dataset may indicate that this estimate may not be well supported.

### Future study & model extensions

Given the bias in juvenile survival and the surprising result of higher mean wild adult survival than breeding program adult survival, further studies in survival are needed to improve these estimates.

In addition to age and location type, there are other covariates that are part of the dataset and could be used in future versions of this analysis such as vaccination status and translocation status that may shed further light on survival for this population of pygmy rabbits.

Future monitoring could be more effective if the monitoring intervals were more regular. Highly irregular monitoring intervals meant that we had to model on a daily scale, which made it difficult to estimate annual-scale variation in survival. Variation in survival is an extremely important aspect of pygmy rabbit population dynamics. Furthermore, developing a sampling plan that ensures that animals can be reobserved (i.e., ensuring adequate spatial coverage) is also critical – these and other aspects of monitoring design will be a topic of ongoing discussions with the working group.

---

**Table C1.** Mean and standard errors of estimated coefficients for both survival and detection. Coefficients were additive; adult age and breeding enclosure location type were set to 0 for survival, and breeding enclosure location type and pellet sign detection type were set to 0 for detection probability.

| Coefficient | Mean estimate | Standard error |
| --- | --- | --- |
| Survival intercept ($\alpha_\phi$; breeding adult) | 6.17722528 | 1.451018e-04 |
| Survival age coefficient for juvenile | 1.09217377 | 2.903291e-04 |
| Survival location coefficient for wild | 0.04369967 | 2.220048e-04 |
| Standard deviation of observation day random effect for survival | 0.92483540 | 5.910263e-05 |
| Detection intercept ($\alpha_p$; breeding pellet sign) | -0.16379805 | 1.534014e-04 |
| Detection location coefficient for wild | -1.58823115 | 1.284713e-04 |
| Detection type coefficient for live capture | -0.30846216 | 1.400105e-04 |

**Figure C2.**
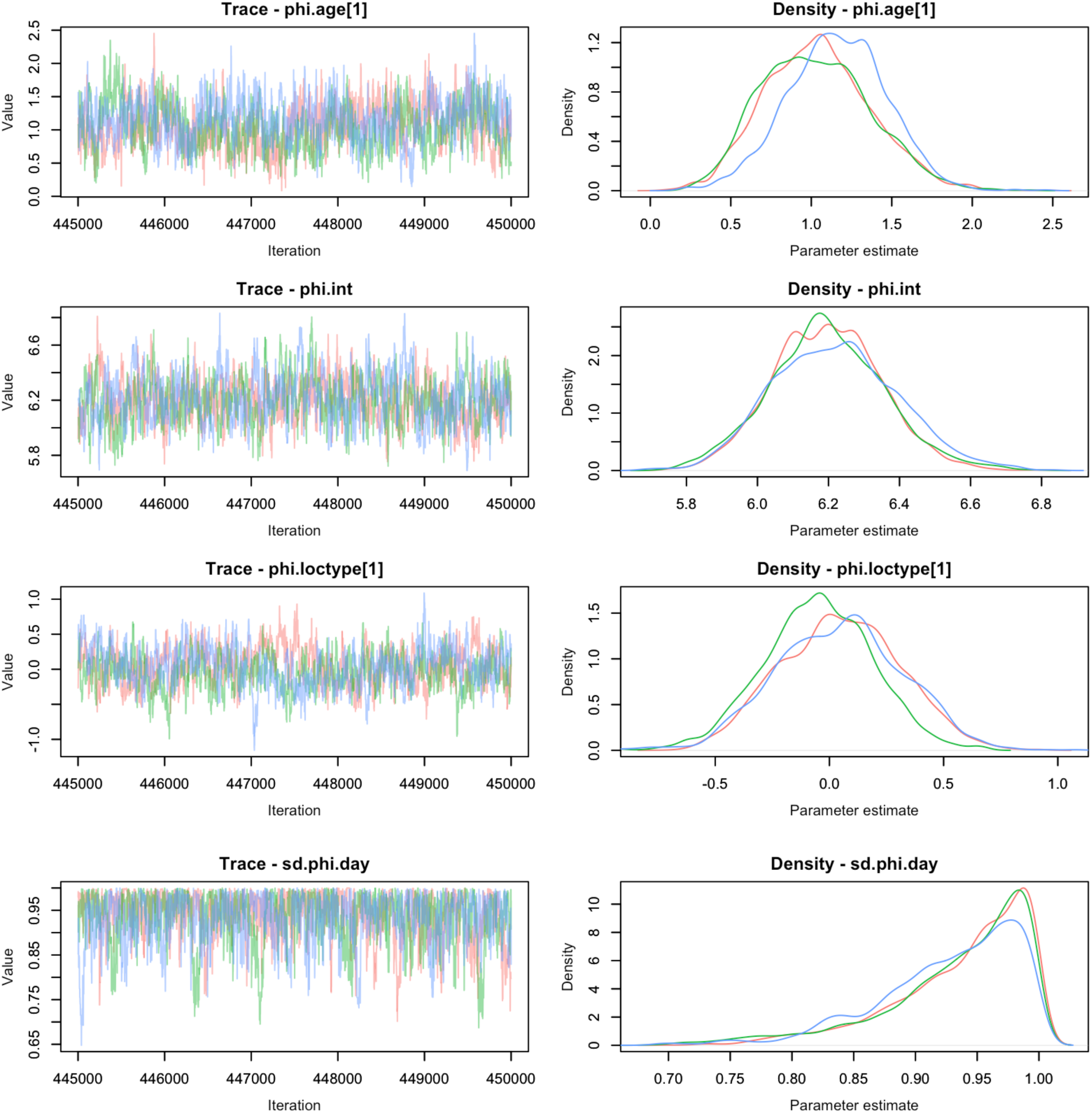
Diagnostic trace plots of the three MCMC chains for each fixed effect coefficient (phi.age[1] is juvenile age, phi.int is the intercept, phi.loctype[1] is wild location type) and the day random effect standard deviation coefficient (sd.phi.day) of survival probability.

**Figure C3.**
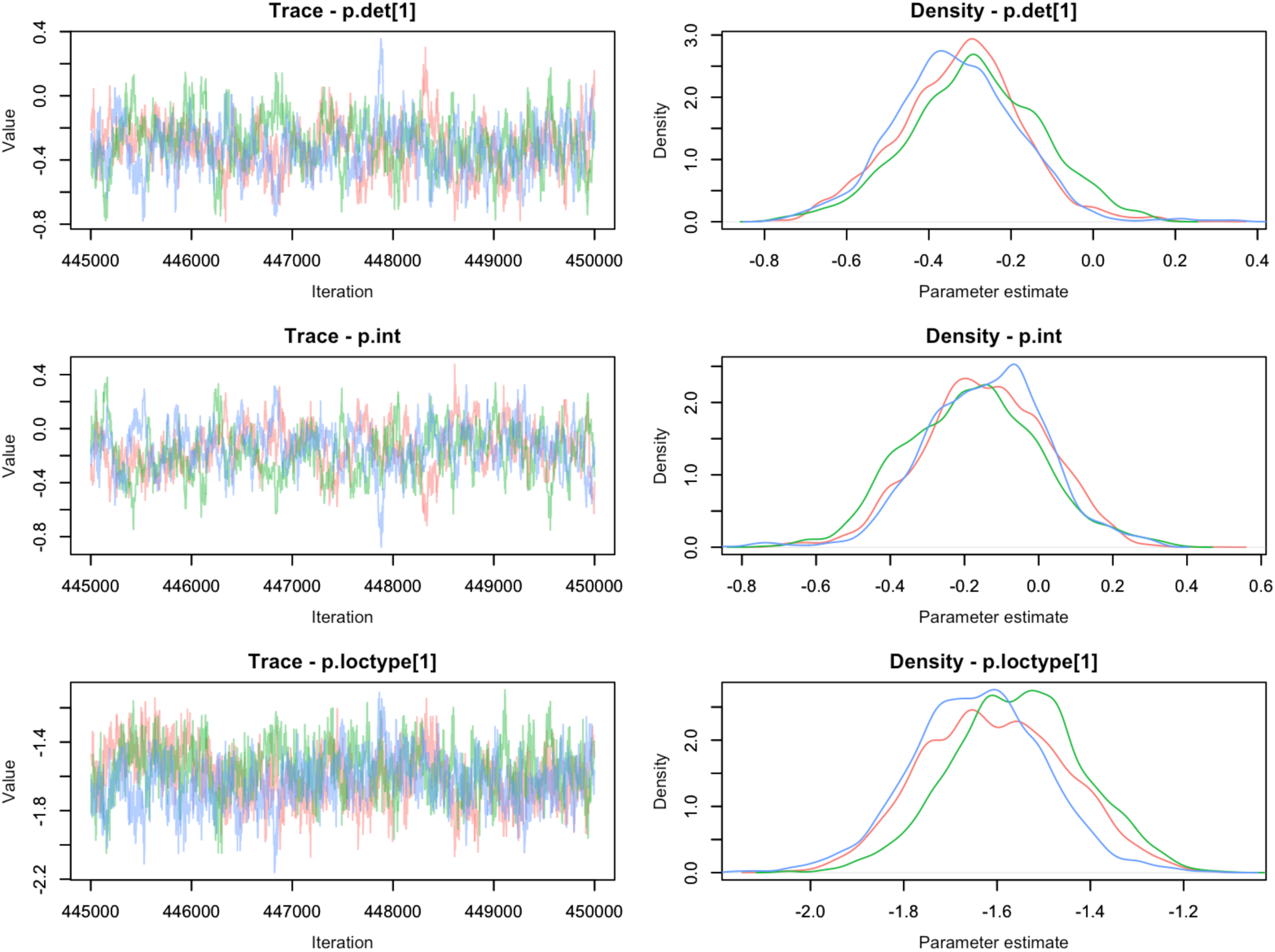
Diagnostic trace plots of the three MCMC chains for each fixed effect coefficient (phi.age[1] is juvenile age, phi.int is the intercept, phi.loctype[1] is wild location type) of encounter probability.

**Figure C4.**
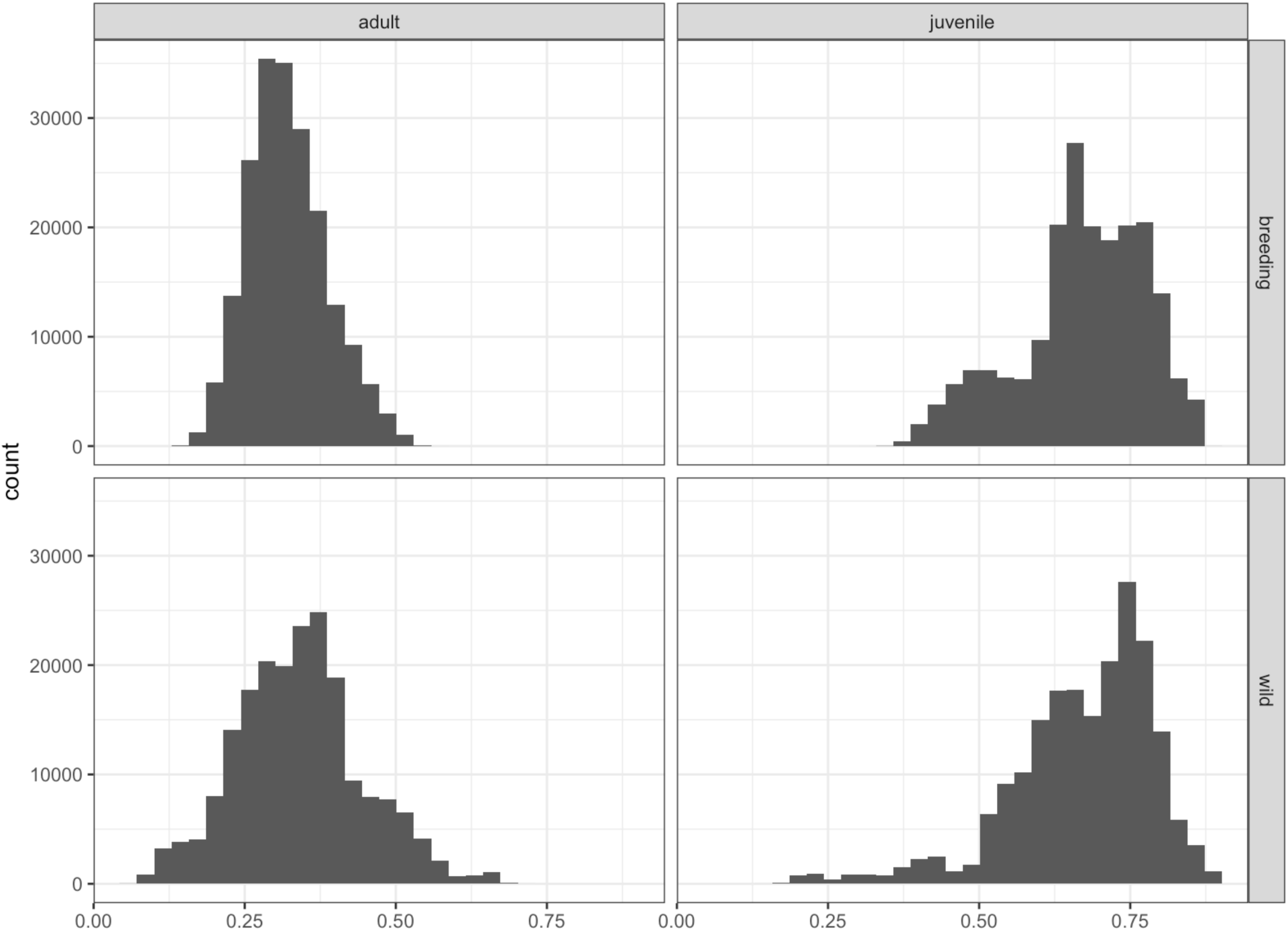
Plots of 200,000 survival probability values by age and location type, calculated with randomly drawn samples of the intercept, fixed effects and random effect standard deviation.

